# Breast cancer stem cells mediated CD8^+^ T cell exhaustion among different molecular subtypes of breast cancer regulated via NOTCH1/RBPJ/PD-L1 axis

**DOI:** 10.64898/2026.05.22.727059

**Authors:** Jasmine Sultana, Aishwarya Guha, Mohona Chakravarti, Goutam Ulgekar, Saurav Bera, Pritha Roy Choudhury, Sukanya Dhar, Juhina Das, Nilanjan Ganguly, Anirban Sarkar, Mariya Saify, Monika Rana, Prodipto Das, Akata Saha, Nirmalya Ganguli, Neyaz Alam, Rathindranath Baral, Anamika Bose, Saptak Banerjee

## Abstract

**Background:** Breast cancer stem cells (BCSCs) contribute significantly to breast cancer (BC) mortality among women globally. It underpins tumor heterogeneity in BC by driving variations in stemness potential and altering immune microenvironment. However, how BCSCs, subpopulations of breast cancer cells from distinct molecular subtypes differentially modulate CD8⁺ T-cell exhaustion and immune dysfunction remain unclear.

**Methods:** We conducted our study from patients with BC of four subtypes. MACS sorted (Lin-CD44+CD24-) BCSCs were prepared for mammosphere formation assay from mastectomies samples. Flow-cytometry was used to analyze breast cancer stem cells (BCSCs). Immunofluorescence, immunohistochemistry, Real Time and Reverse Transcriptase PCR array, Chromatin-immunoprecipitation assay, Transwell, ELISA, Western blotting, Cloning, Transfection, Knockdown, chromatin immunoprecipitation approaches were used to investigate the underlying mechanisms.

**Results:** Here, we report that BCSCs actively participate in tumor progression by modulating effector CD8⁺ T-cells. Triple-negative breast cancer (TNBC), being the subtype with the most adverse outcomes, sustains the enrichment of stem cell regulating transcription factors like NANOG, OCT4 and SOX2 compared to HER2⁺, Luminal B, and Luminal A subtypes. Tumor from TNBC patients exhibited an exhausted phenotype within CD8⁺ T-cell infiltrates with PD1^high^ TIM3^high^ LAG3 ^high^ IFNγ^low^ signature. BCSCs induced increased proportion of exhausted CD8⁺ T-cells, predominantly in the TNBC subtype. Cell-surface Notch1 expression was upregulated in BCSCs across all molecular BC subtypes, with the highest elevation observed in TNBC. Knockdown or inhibition of Notch1 downregulated stemness-associated genes and diminished CSC-mediated induction of CD8⁺ T-cell exhaustion.

Cumulatively, these findings suggest that assessment of high Notch1 and Nanog frequency within BCSCs can guide Notch1-targeted therapies and may formulate for new combinatorial treatment strategies to improve patient outcomes. Additionally, therapeutic targeting of BCSC-intrinsic NOTCH1-NANOG/NOTCH1-PD-L1 axis could represent an effective strategy to reduce stemness programs and alter BCSC-driven CD8⁺ T-cell exhaustion, majorly in aggressive subtypes such as TNBC.

**Conclusions:** BCSCs aggressiveness is perpetuated through Notch1-mediated axis. Targeting Notch1 would reduce stemness (majorly NANOG), survival, as well as prevent CD8⁺ T-cell exhaustion (upregulating PD-1, TIM3, LAG3), thereby weakening tumor progression.

## Background

Heterogeneity within Breast Cancer (BC) emerge from CSCs, influenced by hormone receptor signaling-estrogen receptor (ER), progesterone receptor (PR), and human epidermal growth factor receptor 2 (HER2) (1). BCSC heterogeneity aligns closely with established molecular subtypes (2). Unlike the luminal (ER^+^/PR^+/-^) phenotypes and Her2^+^ subtypes where targeted therapies are available, triple-negative breast cancers (TNBCs), lacking these hormone receptors, display the most diverse BCSC phenotypes, exhibiting poor prognosis (3–5). However, little is known regarding the molecular pathways that modulates functional differences among BCSCs in patients across BC subtypes.

BCSCs are not merely controlled by the TME; rather, through the induction of immune checkpoint ligands such as PD-L1, PD-L2,etc.(6–8). BCSCs actively interact with tumor-infiltrating immune cells-particularly effector CD8⁺ T-cell through modulation of the TME (9). BCSCs derived from TNBC exert pronounced inhibitory effect on CD8⁺ T-cell function despite the presence of immunologically “hot” tumors characterized by abundant tumor-infiltrating lymphocytes (TILs) and elevated PD-L1 expression (10–12). In HER2⁺ BC, BCSC-mediated immune evasion is driven by the AKT/Notch axis and PD-L1/TGF-β signaling (13–15). In contrast, luminal BCSCs induce CD8⁺ T-cell dysfunction primarily through estrogen-driven immune suppression, reduced antigen presentation due to low MHC-I expression, resulting in a chronic, exhausted CD8⁺ T-cell phenotype characterized by elevated PD1, TIM-3, and TOX expression (14,16,17). Despite these subtype-specific observations, how BCSC-driven immune regulation differentially shapes CD8⁺ T-cell exhaustion across all BC subtypes remains unclear. To address this unexplored field, we performed a systematic characterization of BCSC phenotypes across subtypes and evaluated their roles in driving CD8⁺ T-cell dysfunctions.

Here in our present study, we found that BCSCs show distinct enrichment patterns across the four BC subtypes, with variance in stemness regulatory transcription factors associated with aggressiveness. These variant BCSC transcription program had a direct impact on TIL landscape. Furthermore, we are the first to report that subtype specific BCSC (especially TNBC BCSC) were found to be modulating CD8⁺ T-cell dysfunctions via Notch1/Nanog and Notch1/PD-L1/PD1 signaling axis. Together, we demonstrate that BCSC-driven immunosuppressive signaling shapes CD8⁺ T-cell effector functions in a subtype-specific manner. These findings suggest that targeting Notch1/Nanog or Notch1/PD-L1/PD1 axis within BCSCs using immune checkpoint blockade, or targeting transcription factors as novel therapeutic strategy to limit BCSC expansion and improve antitumor immunity in BC.

## Material and Methods

### Reagents, media and antibodies

DMEM: F12K (1:1) media, MEM, RPMI-1640, fetal bovine serum (FBS) were procured from Hi-Media (Mumbai, India). B-27™ Supplement (50X) were obtained from Thermo Fisher Scientific (MA, USA). Growth factors such as human epidermal growth factor (rEGF) and recombinant human basic fibroblast growth factor (rbFGF) were obtained from Merck-Milipore (Darmstadt, Germany). Magnetic isolation kits for human lineage cell panel for lineage sorting and human CD44, human CD24 sorting microbead kits were procured from Miltenyi Biotec (Bergisch Gladbach, Germany). DAPT, Heparin, Fluoroshield™ with DAPI and β-mercaptoethanol were purchased from Sigma (Missouri, USA). Complete list of antibodies is specified in the (**Additional File 2: Supplementary Table –T1).**

### Human solid tumors

Tumors (breast, n=95, including all subtypes) and tumor-adjacent tissues were obtained from patients’ post-surgery, with established norms for histopathology from Chittaranjan National Cancer Institute (CNCI), Kolkata, India after receiving clearance from Institutional Ethical Committee (IEC) (Approval no: CNCI-IEC-SB-20), registered under Central Drugs Standard Control Organization (CDSCO), Govt. of India. For breast carcinoma, TNM staging grade FIGO classifications are followed. Detail information is provided in (**Additional File 2: Supplementary Table –T2)**.

### Data Availability

Data analysis from patient records was performed for individuals diagnosed with BC at Chittaranjan National Cancer Institute, Kolkata following approval of Institutional Ethics Committee (Approval no: CNCI-IEC-SB-20) dated 25.04.2022. Luminal A, Luminal B, Her2^+^ and TNBC patients at stages I–IV were considered for inclusion. All data analyzed during this study are available from the corresponding author on request.

## Inclusion criteria for patients

Patients were eligible after qualifying following criteria:

(a) Patients with cytologically or histopathologically confirmed malignant Luminal A, Luminal B, Her2^+^ and TNBC tumors, irrespective of stage and grade.
(b) Female patients within the age group of (25–70) years.
(c) Patients with available complete clinical records detailing disease recurrence status,

## Exclusion criteria for patients

Patients were excluded after obtaining following criteria:

(a) Patients undergoing Neoadjuvant Chemotherapy (NACT) treated samples,
(b) Male patients with breast tumors.

### Tumor tissue processing for cell-culture, flow-cytometry and cryo-sectioning

Single cell suspension was aseptically generated from excised solid human tumor specimens after mechanical dissociation and subsequent digestion with collagenase IV, rinsed in PBS, comprising 2mM penicillin-streptomycin. The cell residue was strained through 70µM cell strainer and centrifuged at 1800 RPM for 5 mins for single cell suspension. The cells were used for flow-cytometric staining. For cryo-sectioning, solid tumors were fixed in 4% paraformaldehyde at RT for 2h followed by incubation in 30% sucrose solution at 4°C overnight. Lastly, they were kept at –80°C till cryo-sectioning after being rapidly frozen in liquid nitrogen.

### Cryo-sectioning

Fixed tissue samples were embedded in an optimal cutting temperature (OCT) compound (Leica Biosystems, Wetzlar, Germany). These samples were collected on poly-L-lysine-coated slides, sectioned (5µM) on a Cryostat (Leica CM1950, Wetzlar, Germany, RRID: SCR_018061) and stored at –80°C until needed for further use.

### Histology, hematoxylin-eosin (HE) staining and quantification

Cryo-sectioned tissues were PBS-rinsed and were stained with HE ensuing standard staining protocol. Their quantification was accomplished by using ImageJ software RRID: SCR_003070.

### Purification of BCSCs and CD8⁺ T-cells by magnetic cell sorting from tumor cells

Tumor cell suspensions were magnetically sorted with CD44/CD24/Lineage-cocktail antibodies as per requirement. Cell purification was performed according to the manufacture’s protocol (MicroBead kit, Miltenyi Biotech, Germany). Lineage positive cells were purified for CD8^+^ T-cells and Lin-CD24^-^CD44^+^ for CSC population. The purity of sorted cells was checked using flow-cytometry.

### BCSC enrichment milieu

Single cell suspension from tumor cells were magnetically sorted for BCSC and prepared in BCSC enrichment media containing serum-free DMEM: F12K (1:1) with 1% B-27™ supplement (50X) and freshly enhanced with human-rEGF (20 ng/mL), human-rbFGF (20 ng/mL), heparin (40 ng/mL). They were cultured on ultra-low adherent plate (Corning, New York, USA) and incubated at 37°C with 5% CO_2_ for six days and supplemented with fresh media at every three days. After six days, the tumorspheres formed were dissociated with mild trypsinization and centrifuged for single cell suspension for further studies. These single cells were called as primary mammosphere. For secondary mammosphere formation, similar protocol was followed after enrichment of single cell suspension formed from primary mammosphere. The generated tumorspheres were counted from random fields and their area was calculated using ImageJ software.

### Co-culture assay

Sorted Lin^+^ cell subtypes were cultured in BCSC enrichment media as described earlier and with RPMI-1640 complete media (1:1) containing βME (beta mercapto ethanol) with sorted BCSCs from each subtype in a 10:1 ratio for 48 h. After incubation period, CD8⁺ T-cell suspension was sorted and centrifuged for further downstream assays.

### Reverse transcriptase (RT) PCR

RNA content was extracted using Trizol (Ambion, Thermo Fisher Scientific, MA, USA) and respective cDNA was extracted using Revert Aid First Strand cDNA Synthesis Kit (Thermo Fisher Scientific, MA, USA) according to the manufacturer’s protocol. RT-PCR was done using 2X Go Taq Green Mix (Promega, WI, USA). Electrophoresis was carried out using 1.5% agarose followed by staining with ethidium bromide. PCR products were visualized on ChemiDoc XRS+ (BioRad Laboratories, CA, and USA) and band intensity was quantified by using ImageJ software (RRID: SCR_003070). Gene specific primers that are used in PCR are listed in (**Additional File 2: Supplementary Table –T3).**

### Immunofluorescence

Samples or sections, targeted for immunofluorescence staining were washed with 1X PBS. Cells were fixed, perforated with 0.15% Triton X-100 prior to blocking with 5% BSA at 4°C over-night for staining intracellular molecules (otherwise avoided for only extracellular staining) (18). Primary antibodies (1:200 dilution) for specific antigen were added to the section and incubated over-night at 4°C. Fluorescent-dye-conjugated secondary antibodies (1:200 dilution) were added (if primary Ab is not tagged) and incubated for 2.5 h at RT. Samples were washed properly and mounted on slides (for mammosphere), with Fluoroshield DAPI (Abcam, Cambridge, UK). Images were taken with Olympus-BX53 (Olympus Lifesciences, Tokyo, Japan) (RRID: SCR_022568) and Confocal Microscope (FV4000, Olympus, Japan) (RRID: SCR_022568). Fluorescence intensity was measured using Image-J software and corrected total cell fluorescence (CTCF) was evaluated by using the formula {CTCF = Integrated density− (Area of selected cell × Mean fluorescence of background readings)} (19).

### Extraction of cytosolic and nuclear proteins

Ice cold cytoplasmic extraction buffer was added to the pellet of sample of interest, vortexed properly and incubated at 4°C for 1 h. The cells were centrifuged at 8000 g for 15 mins and the supernatant was collected as cytosolic fraction. The pellet was then dissolved with nuclear extraction buffer with proper vortexing and incubated at 4°C for 30 mins. The samples were centrifuged at 14000g for 30 mins. The supernatant thus acquired was the nuclear fraction. Using Bradford assay, protein concentrations of the lysates were determined.

### Western Blotting

Western blotting was executed as specified. Targeted cellular protein lysates were mixed with Laemmli buffer (50 µg/ ml) and boiled for 15 mins. 25-30μg of the protein lysates was fractionated by 12% SDS-PAGE and transferred onto nitrocellulose membrane using BioRad Gel Transfer system followed by blocking with 7-8% BSA for 3 h. The bands were probed with definite HRP-conjugated secondary antibodies (1:10,000 dilution) for 2hrs after incubating overnight with specific primary antibodies (1:500 dilution) at 4°C. The bands were developed using ECL Kit (Advansta, CA, USA). Band intensities were quantified using Image Lab 6.2 software (Bio-Rad, California, USA).

### Immunohistochemistry

Desired tissue sections were harvested initially on glass slides coated with poly-L-lysine, washed in 1X PBS for OCT removal. In order to block endogenous peroxidase, sections were treated with 3% H_2_O_2_ (Merck 17544) in methanol for 30 mins. Blocking was carried out after incubation with 5%BSA for 30 mins in order to block non-specific interaction subsequent to incubation with primary antibody (1:150 dilution) overnight. HRP tagged secondary antibody (1:1000 dilution) was added to the sections post comprehensive washing with PBS-Tween 20. AEC Substrate (Vector Laboratory SK4200) was applied to develop chromogenic color according to manufacturer’s protocol. Counterstaining was accomplished using Hematoxylin (Merck-HX68597049) for 40-60 secs and then mounted with Vectormount (Vector Laboratories H5501). Image was procured using the Carl Zeiss Plan Achromat bright field microscope along with the Axiocam color camera. Image analysis was conducted by Image J IHC plugin software (20).

### Transwell assay

CD8⁺ T cell population was co-cultured with BCSC of Luminal A/Luminal B/Her^+^/TNBC cells in a 10:1 ratio for 48 h. in the presence of a 0.4 µM transwell membrane (Hi-Media, Mumbai, India). After incubation, membrane was removed, and the cells were examined for further downstream studies.

### Cell lines and cultures

MDAMB-231 (RRID: CVCL_0062) cell line was acquired from National Centre for Cell Sciences (NCCS), Pune, India. Authentication of the cell line was conducted via STR profiling by cell origin. Cells were proliferated and stored according to the provider’s directions. Above cell line was cultured in complete media added with 10% (v/v) heat inactivated FBS, 2mM L-glutamine, 100 U/ml penicillin and 100μg/ml streptomycin in 5% CO_2_ in incubator at 37° C. Monitoring of cell lines were regularly conducted for perfect cell morphology and probable mycoplasma contamination. If mycoplasma contamination detected, it was removed using EZkill^TM^ Mycoplasma Elimination Kit (Hi-Media, Mumbai, India). Cells were stored after two passages and experiments were processed within 10-12 passages.

## Tumor antigen-loaded DCs generation and co-culture assay

DCs were generated from human PBMCs (21). Monocytes derived from PBMCs were selectively adhered in 90mm petri-plate. The adhered monocytes were then cultured with GM-CSF (800 U/mL) and rIL-4(500 U/mL) in AIM-V media with 10%FBS until they differentiated into immature-DCs for seven days of culture. For, DC maturation and tumor-antigen loading, immature DCs were incubated in AIM-V media with 10% FBS with rGM-CSF (800 U/mL) and rIL-4 (500 U/mL), rTNF-α (5ng/mL), rIL-1β (5 ng/mL), rIL-6 (150 ng/mL) and tumor-antigen (5 µg/mL) overnight. Later, mature antigen loaded DCs were collected with gentle tapping. CD8^+^ T-cells, isolated from PBMC of healthy donors were incubated with Tu-Antigen-loaded DCs (30:1 ratio) for 2 days. Then these activated CD8⁺ T-cells were cultured with sorted BCSCs from each subtype in a (10:1) ratio for 48 h. For further studies CD8⁺ T-cells were sorted out.

### Flow-cytometry

Intended cells were counted and stained with antigen specific fluorescent-tagged antibodies (1:200 dilution) at 4°C for 30 mins in the dark. For intracellular molecules staining, cells were treated with permeabilization buffer (0.3% saponin in PBS). Targeted cells were finally washed with 1xPBS and fixed with 1% paraformaldehyde. Data was acquired using BD LSR Fortessa X-20 Cell Analyzer (RRID: SCR_018655) (Becton Dickinson, New Jersey, USA). For data retrieval Cell Quest Pro 5.1 and for data analysis FlowJo (10.8.1) (RRID: SCR_008520) (Becton Dickinson, New Jersey, USA) (RRID: SCR_018655) were used. Gating strategies are provided in the main and supplementary figures, and representations of data were made as per convenience of explanation.

### ELISA

From test groups, antigens were collected and were anchored on 96-well microtiter plates. Cells were blocked with 5% BSA at RT for 30 mins. Specific primary antibody (1:500 in 1% BSA) (50 µl/well) was added overnight at 4°C. The respective wells were washed 2-3 times with PBS-Tween 20 (200 µl/well), followed by incubation (3 h in dark) with HRP-conjugated secondary antibody (1:1000 in 1% BSA) (50 µl/well). Finally, TMB substrate (BD OptEIA, BD Biosciences) was added in these wells resulting a visible colored-product (light blue/cyan). Later this color was changed into yellow after terminating the reaction with 1 (N) H_2_SO_4_. Optical density was measured at 450 nm by Spectramax i3X (Molecular Devices, San Jose, USA) (RRID: SCR_026346). The result was quantified by SoftMax Pro 7.1 software (RRID: SCR_014240).

### Protein-protein interaction visualization

String (Search Tool for the Retrieval of Interacting Genes/Proteins) database version 8.0 RRID: SCR_005223 was utilized to unravel the interactions between numerous proteins of interest. An interactome map presenting the interrelationship among proteins was obtained based upon direct and indirect interactions. Multicolor lines depict the relationship among proteins. The network properties include nodes (the number of proteins), edges (connections), node degree (average interactions) and clustering coefficient, which indicates varieties of interactions among them.

## Chromatin-immunoprecipitation assay

Two thousand nucleotide sequence of Nanog and Pd-l1 of Homo sapiens were probed in the eukaryotic promoter database (EPD) and transcription binding motifs (TFBM) for binding to RBPJ were identified using JASPAR CORE (RRID: 309 SCR_003030). Then we found consensus sequence within TFBM sites for nanog and Pdl-1 through transcription factor binding site database (TRANSFAC 8.3) (RRID: SCR_001624), applying the matrix-based TFBM-prediction algorithm named PROMO in the ALGGEN (RRID: SCR_016926) (BarcelonaTech) web server. Generated consensus sequences were aligned with the corresponding TFBMs containing short stretch of DNA in the sequence alignment editor, BioEdit 7.2.5 (NCSU, North Carolina, USA), to locate the exact TFBM regions in those genes (–658th to –650th nucleotides for Rbpj binding in Pdl-1 gene sequence, along with –955th to –939th nucleotides for Rbpj binding in the Nanog gene). ChIP assay was conducted according to the manufacture’s protocol (Millipore, Darmstadt, Germany). Cultured cells (1×10^6^) per group were fixed with 1% paraformaldehyde. These cells were washed with chilled PBS (containing 1 mM PMSF and 1 µg/ml pepstatin A as protease inhibitors) and lysed within SDS-lysis buffer by a 10 min’ incubation on ice. The DNA content was sheared to small stretches by ultrasonication (5 pulses of 30%, 5 s each in UP400S, Ultrasonicator, Hielscher, NJ, USA) (22,23). The lysate formed was cleared using Salmon Sperm DNA/Protein A Agarose-50% Slurry followed by agitation at 4 °C overnight and brief centrifugation in order to spin down the beads. A certain amount of supernatant consisting of all protein-DNA complexes was collected from samples as input DNA (positive control) prior immunoprecipitation. The desired DNA-protein complexes were incubated at 4 °C with primary antibody Rbpj (5µg), with rotation. It was captured using anti-goat IgG-coated agarose beads (Sigma-Aldrich, MO, USA). Eluted DNA was extracted using phenol/ chloroform and precipitated using 70% ethanol. Presence or absence of the respective TFBM (for Rbpj binding near nanog and Pd-l1 gene promoters) among these DNA were identified using qRT-PCR amplification of those particular sites using promoter specific primer-sets for Rbpj binding site for Nanog and Pd-l1 promoters. The primers used for Chromatin Immunoprecipitation (ChIP) are added in the **(Additional File 2: Supplementary Table-T3)**.

### Statistical analyses

The exact numbers of experimental data (n values) are mentioned in the respective figure legends. All results represent the mean±SD of independent repeats. Statistical significance was recognized from Two-tailed unpaired Student t-test (for two groups) or One-way ANOVA, followed by Tukey’s multiple comparisons test (for more than 2 groups) for ungrouped data. While for grouped data, Two-way ANOVA (more than 2 groups) was used, followed by Tukey’s multiple comparisons test. Positive or negative correlation was inferred from Pearson Correlation co-efficient. All statistical analysis and heatmaps were generated by Graphpad Prism 8.4.2 software (GraphPad Software, San Diego, CA, USA) (RRID: SCR_002798). Experimental groups achieving a p ≤ 0.05 were measured as significant.

## Methodology for Cloning, transfection and knockdown

### Cloning of Notch1 shRNA constructs

Four different short hairpin RNA (shRNA) sequences targeting human Notch1 (shRNA1–4) were designed as complementary oligonucleotides **(Additional File 2: Supplementary Table – T4)**. For annealing, forward and reverse oligos (8 μL each) were mixed with 2 μL of NEB 2.1 buffer and incubated at 98°C for 10 mins. The mixture was allowed to cool gradually in a water bath to enable proper annealing of the strands.

The cloning strategy was designed in-silico using snapgene software (Dotmatics, snapgene, USA). The detailed cloning strategy is mentioned in (**Additional File 1: Supplementary Figure S5).** Briefly, pRNAT CMV 3.1-EGFP-Neo vector (Genscript, Nanjing, China) was digested with BamHI and AflII restriction enzymes (New England Biolabs, USA) to generate compatible staggered ends and subsequently gel-purified. Annealed oligos were ligated into this digested vector using T4 DNA ligase. The ligation mixture was transformed into competent E. coli DH5α cells and plated on LB agar plates containing ampicillin (100μg/ml). Additional mapping was performed with SalI and EcoRV (New England Biolabs, USA) to further confirm the integration. Plasmids showing expected digestion patterns were sent for Sanger sequencing to validate correct orientation and sequence integrity **(Additional File 1: Supplementary Figure S6)**. Based on preliminary screening in MDA-MB-231, we have selected shRNA2 for stable cell generation. To this end, we cloned Notch1 shRNA2 in pCMV-EGFP miR-E backbone **(Additional File 1: Supplementary Figure S8 and S9)**, followed by subcloning into pLJM1-EGFP (Addgene#19319) vector **(Additional File 1: Supplementary Figure S11 and S12)** to generate pLJM1-EGFP-Notch1-shRNA2 for lentivirus mediated transduction (24).

### Cell culture, transfection and generation of stable Notch1 knock down MDAMB-231 cells

MDAMB-231 cells were maintained in Dulbecco’s Modified Eagle Medium (DMEM) supplemented with 10% FBS and antibiotics-antimycotic at 37 °C in a 5% CO₂ atmosphere. For transfection, cells were seeded at densities ranging from 2.5 × 10⁵ to 3.0 × 10⁵ cells/well in 6-well plates. After 48 h of incubation, cells were transfected using FA6-bPEI reagent, with plasmid DNA (1.5μg per well) at a DNA: FA6-PEI ratio of 1:1. Both the untransfected controls and vector-only controls were included in each experimental set. The transfected cells were checked by using BD FACSMelody Cell Sorter (BD Biosciences, USA): RRID:SCR_023209. To generate lentivirus particles, we co-transfected HEK293T cells with pMD2.G, psPAX and pLJM1-EGFP-Notch1-shRNA2 vectors and spent media containing lentivirus particles was harvested after 72 h. For transduction, MDAMB-231 was seeded in T25 flask, followed by treatment with lentiviral spent media diluted in growth media at 1:1 ratio for 24 h. We observed EGFP positive cells after 24 h culture, which was then subjected to puromycin selection to generate stable cell line (25).

### Fluorescence microscopy

Fluorescence imaging was performed 24 h and 48 h post-transfection/post-transduction using an inverted fluorescence microscope with brightfield and EGFP channels were acquired in parallel **(Additional File 1: Supplementary Figure S7 and S10)**. EGFP expression was used as a marker of transfection efficiency, while brightfield imaging was used to assess cell morphology and viability. Images were visualized using ZOE Fluorescent Cell Imager (Bio-Rad, USA)-RRID:SCR_019973.

### RNA isolation and cDNA synthesis

Cells were harvested at 24-48 h post-transfection or from confluent stable Notch1^KD^ MDAMB-231 for RNA extraction using trizol reagent. RNA pellets were air-dried, resuspended, and quantified spectrophotometrically. For cDNA synthesis, 500–1000 ng of total RNA was reverse transcribed using either random hexamers or oligo (dT) primers in technical replicates.

### PCR and quantitative RT-PCR

PCR amplification of notch, actb, and gapdh was performed using specific primers **(Additional File 2: Supplementary Table – T5)**. C-1000 Thermal Cycler (Bio-Rad, USA): RRID:SCR_019688 was to perform qRT-PCR. PCR products were resolved on agarose gels to confirm expected band sizes. The densitometry was performed using the ImageJ software. For qRT-PCR, SYBR Green chemistry was used with 40 cycles of denaturation and annealing/extension. Melt-curve analysis was performed to identify primer-dimers or nonspecific amplification. actb and gapdh were used as internal controls for normalization. To evaluate the functional effects of Notch1 knockdown, expression of nanog, rbpjκ, pdl1, and hes1 was analyzed by conventional PCR using gene-specific primers **(Additional File 2: Supplementary Table – T5)** in selected sets of transfected cells. These genes were selected due to their established roles as downstream targets of Notch1 signaling. Images of PCR products were taken by ChemiDoc MP Imaging System (Bio-Rad, USA):RRID:SCR_019037.

### Statistical analysis

All the statistical analysis was performed using the graphpad prism software. The data (Mean ± SEM) is indicative of n= 4 independent experiments and analyzed by two-tailed paired Student t-test. All statistical analysis was generated by GraphPad Prism 8.4.2 software (GraphPad Software, San Diego, CA, USA). Experimental groups achieving a p ≤ 0.05 were measured as significant.

## Results

### Breast cancer stem cells (CD44^+^CD24^-/low^) and Nanog are differentially distributed within molecular sub-types of breast cancer

Heterogeneity of BC varies across all the subtypes at both genetic and morphological levels. To investigate the distribution and molecular underpinnings of stemness-regulating factors driving their distinct aggressiveness, the BCSC populations were isolated from the four molecular subtypes of BC patients without any prior treatment (NACT-Neoadjuvant chemotherapy). Single cells from the breast tumor-biopsies (n=11, from each subtype) were isolated and sorted based on the Lin^-^CD44^+^CD24^-^ surface phenotype (Fig. 1A).

**Figure 1:**
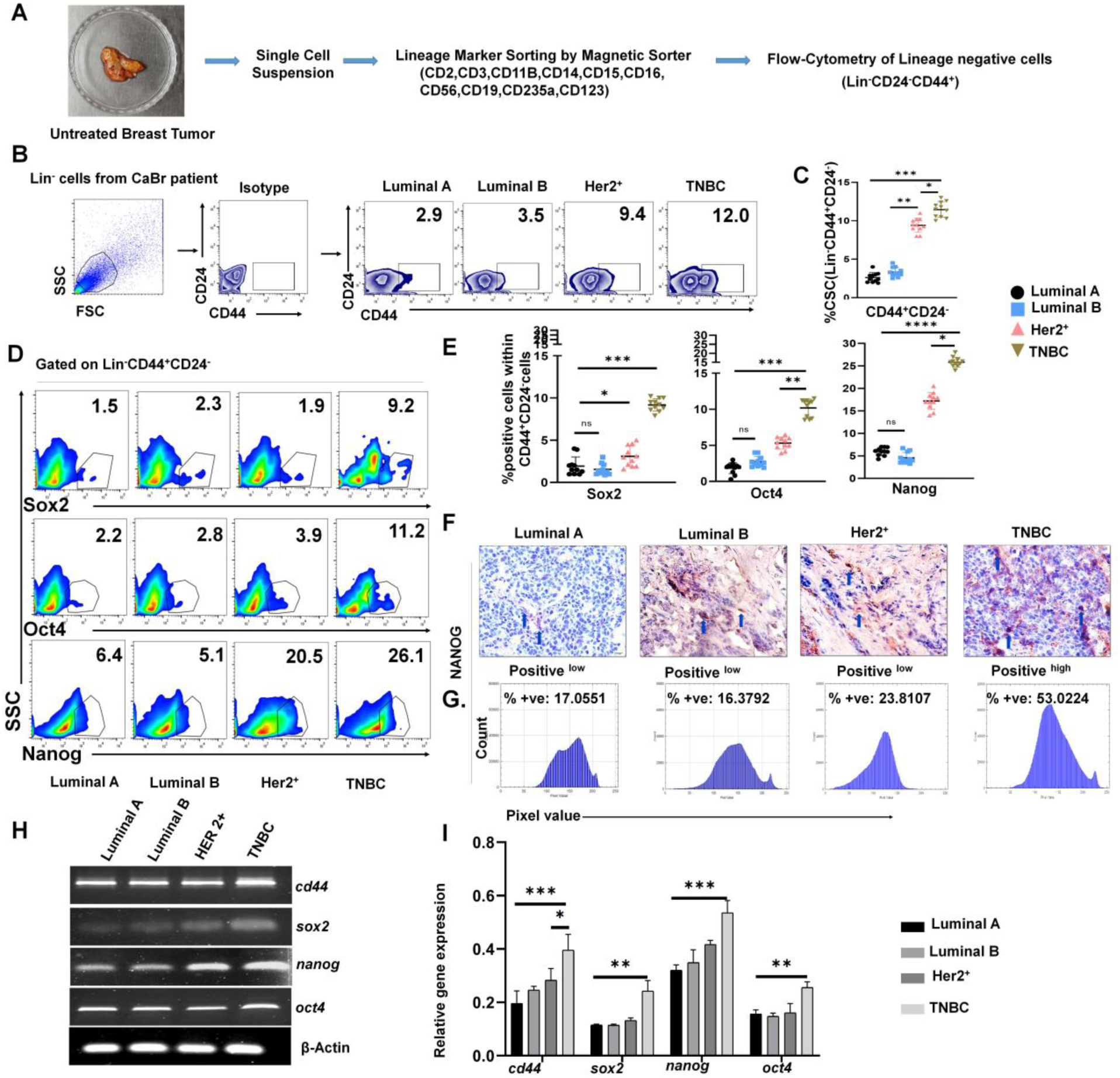
CD44^+^ CD24^−/low^ cells in BCSC correlates positively with Nanog frequency within TME of BC subtypes in pre-enrichment-setting. (A) Schematic representation of isolation of lineage negative population from breast tissues isolated from BC patients. (B) Representative flow-cytometric plots for population (Lin^-^CD44^+^CD24^-^) in BC biopsy samples. (C) Scatterplots in each case demonstrating frequency of individual populations with mean±SD in all subtypes (n=11). (D) Representative flow-cytometric-plots characterizing status of stem-cell-regulatory factors – SOX2, OCT4 and NANOG (gated within CD44^+^CD24^-^) from BC patients. (E) Scatterplots depicting frequency of these transcription factors of individual breast subtypes (n=11 in each cohort) with mean±SD, analysis was conducted by One-way ANOVA followed by Tukey’s multiple-comparison test. (F) Representative IHC of NANOG expression in tissue sections from Luminal A, Luminal B, HER2-enriched, and TNBC subtypes. (G) IHC Profiler plugin assessed with ImageJ displayed markedly elevated nuclear NANOG levels compared to other subtypes at 40 X magnification. (H) Status of *sox2, nanog* and *oct4* within CSC population across all subtypes via RT-PCR. (I) Relative fold changes in gene expression are presented by bar-graphs (mean± SD); p value drawn from repeated measures two-way ANOVA. (n=5). (p < 0.05). (C, E, G) *\*, p < 0.05; **, p < 0.01; ***, p < 0.001; ****, p < 0.0001*; *ns, nonsignificant* are indicated.

Tumor-derived sorted BCSCs were subjected to flow-cytometry to quantify and assess expression of stemness-associated transcriptional factors. The mean proportion of the BCSC subpopulations was found to be highest within TNBC tumors (11.45% of BCSC) than luminal type tumors (2.62 % and 3.33 % of BCSC in luminal A and B respectively). However, in Her2^+^ BC the percentage elevation of BCSCs were intermediate (9.41% of BCSC) (Fig. 1B, 1C). To illustrate the underlying transcriptional architecture governing stemness, we profiled the expression of key pluripotency-associated transcription factors. Notably, TNBC CSCs demonstrated the highest frequency of NANOG (25.21%) compared to luminal (5.78 % and 4.48% in Luminal A and B respectively) and Her2^+^ BC (16.60%). Additionally, OCT4 and SOX2 expression levels were also found to be elevated in TNBC samples, while in HER2^+^ and luminal subtypes, comparatively lower CSC abundance and reduced expression of associated markers were observed (Fig. 1D, 1E). Cumulatively, this study suggested that BCSC phenotype showed a significant relationship with TNBC population as this subtype are highly enriched with pluripotent transcription factors OCT4, SOX2 and NANOG.

In line with high Nanog expression in protein levels, immunohistochemistry on breast tumor tissue also demonstrated high NANOG expression in TNBC tissues, compared to other subtypes. Immunostaining intensity was analyzed, which revealed low positive staining (+), in Luminal A and Luminal B tumors indicating minimal NANOG expression. HER2-enriched tumors exhibited intermediate (++) to high (+++) NANOG expression. In contrast, TNBC tissues showed a noticeable increase in low (+) positive to high (++++) positive nuclear staining, indicating significantly higher levels of NANOG expression compared to the other subtypes (Fig. 1F). This significant surge in NANOG percentage validates the probable role of NANOG in the altered BCSCs fate (Fig. 1G).

Quantification of genes involved in CSC stemness by RT-PCR based analysis (*cd44*, *sox2, nanog* and *oct4*) revealed that the expression of *sox2, nanog* and *oct4* was also significantly upregulated in CSCs derived from TNBC followed by Her2^+^>LumB>LumA. Similar in line with this report, it was observed that TNBC exhibited higher expression of these genes as compared to their counterparts attributing to their aggressive state (Fig. 1H, 1I).

## Differential expression of Nanog and other stemness markers in secondary enrichment culture positively correlated with the tumorigenic potential of CSC

Following the inherent characterization of tumor-sorted BCSCs within different molecular subtypes of BC, we explored the tumorigenic potential of BCSCs *in-vitro* across serial passages. Persistence of tumorigenicity with passages functionally validates stemness (26).

Lin^-^CD44^+^CD24^-^ sorted cells are considered as stem-like but enriching them in CSC specific enrichment media promotes the expansion of BCSCs and maintains the stability of that phenotype. For this purpose, single cells obtained from four different molecular subtypes of breast tumor were magnetically sorted to acquire BCSCs. Cells were propagated for 7 days in a serum-free conditions and this culture period culminated in the formation of primary mammospheres (Fig. 2A) and were further dissociated into single cells in enrichment media for secondary mammosphere formation (Fig. 2B).

**Figure 2:**
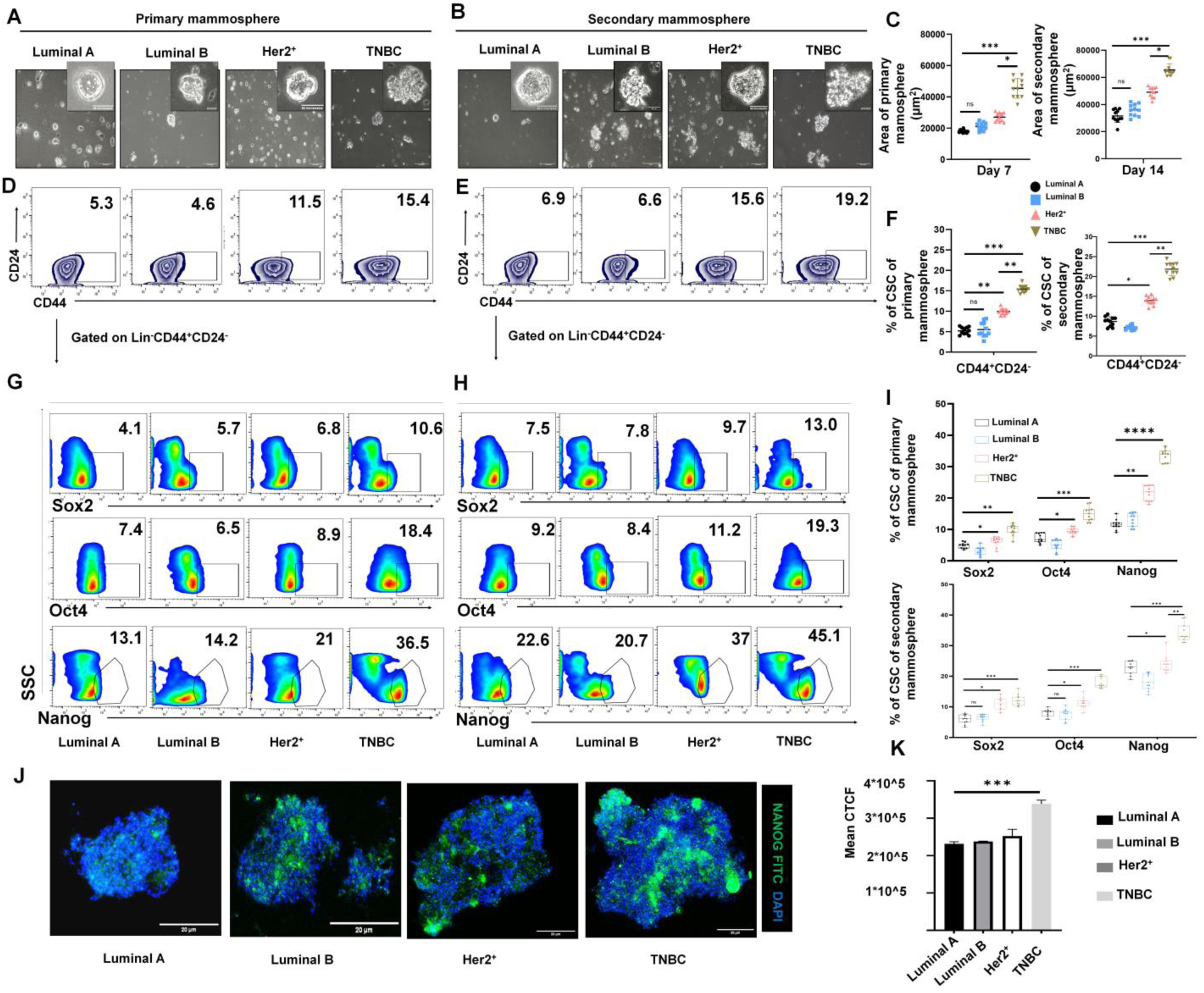
CD44^+^CD24^−/low^ BCSCs and Nanog frequency generated from post-enrichment setting are significantly greater than pre-enrichment one. (A, B) Representative images of primary and secondary tumorspheres for BCSCs magnetically sorted (Lin^-^CD44^+^CD24^-^) from Luminal A, B Her2^+^ and TNBC patient samples and cultured for tumorsphere assay (10X magnification); insert in 20X. (C) Represents scatter plot, (mean±SD) for tumorsphere area of primary and secondary mammosphere of all subtypes (n=11). (D, E) Flow-cytometric zebra plots signifying changes in CSC-frequency in four subtypes with or without enrichment in primary and secondary mammosphere. (F) Scatter plots representing CSC frequency of primary and secondary mammospheres of four subtypes. (G, H) Representative pseudo-color flow-cytometric plots demonstrating variations in the frequencies of transcription factors regulating stem cell fate – OCT4, SOX2, NANOG within BCSC population of respective groups (n=11) in primary and secondary mammosphere. (I) Box and whiskers representing transcription factors frequency across Luminal A, Luminal B, Her2^+^and TNBC tumor samples of primary and secondary mammosphere. Statistical interpretation based on one-way ANOVA followed by Tukey’s multiple-comparison test (n=11). (J) Representative immune-fluorescence micrographs at 100X magnification of mammosphere from BCSCs of four subtypes stained with NANOG-FITC. (K) Cytofluoromicrographs and Mander’s co-localization co-efficient (M1) for NANOG and nuclear signal co-localization are provided in lower panel. (C, F, I) *\*, p < 0.05; **, p < 0.01; ***, p < 0.001; ****, p < 0.0001*; *ns, nonsignificant* are indicated.

In mammosphere assays, primary mammosphere depicts the abundance of stem cells, while secondary mammospheres portrays the intrinsic self-renewal capacity of the stem cells (27). In these anchorage-independent conditions the spheroids from four different subtypes displayed different tumorsphere counts, with the highest count observed in case of TNBC (Additional File 1: Supplementary Fig. S1-A). To delineate the transcriptional circuit nourishing CSC enrichment, BCSCs throughout primary (after 7 days) to secondary mammospheres (after 14 days) across all the four subtypes of BC were analyzed. mRNA expression profiling by RT-PCR was employed to investigate the association of the stemness-related transcription factors *oct4*, *sox2*, and *nanog* (Additional File 1: Supplementary Fig. S1-B, C). From primary (1^0^) to secondary (2^0^) mammosphere, a stepwise elevation in the expression of these transcription factors was witnessed. Flow-cytometry results revealed a prominent increase in BCSC percentage following culture in enrichment media. The rate of secondary sphere formation was remarkably greater than that of primary mammosphere and area of mammospheres formed per well is slowly increased from day 7 to day 14 (Fig. 2C). Among these, TNBC exhibited the most robust expansion of BCSC population (1^0^ vs 2^0^ is 15.45% vs 21.85%) than Her2^+^ (1^0^ vs 2^0^ is 9.96% vs 13.96%). Luminal A (1^0^ vs 2^0^ is 5.21% vs 8.68%) and Luminal B (1^0^ vs 2^0^ is 5.5% vs 7.16%) BC harbored the least enriched population of BCSCs (Fig. 2D, 2E, 2F).

Following the quantification of BCSC frequency within the mammospheres, the investigation was next directed towards evaluating the expression of the core stemness regulators within these cellular subsets. The expression of OCT4, SOX2 and NANOG within BCSCs in BC patients (*n* = 11 in each subtype) were explored (Fig. 2G, 2H, 2I). However, NANOG emerged as the most consistently up regulated across all four subtypes, even in BCSC poor Luminal A and Luminal B tumors. No prominent changes were noted in SOX2 level where as OCT4 level were slightly elevated in Her2^+^ and TNBC subtypes of BCSC (Fig. 2G, 2H, 2I). Whereas, TNBC BCSCs displayed the highest induction of all three transcription factors, consistent with their aggressiveness. Confocal microscopy confirmed augmented Nanog nuclear accumulation distinguishing BCSCs across all subtypes; however, TNBC displayed maximal expression and highlighting its dominant stemness-driven phenotype (Fig. 2J, 2K).

## Intra-tumoral CD8^+^ T-cells from different BC subtypes showed distinct functional profile and T cell exhaustion positively correlated with disease aggression

Apart from intrinsic cancer cell stemness, functional impairment of infiltrating CD8⁺ T-cells also plays a distinctive role in tumor progression and prognosis of BC. To seek deeper understanding about the CD8⁺ T-cell status, tumor infiltrating CD8⁺ T-cells were sorted from post-operated samples (Luminal A, Luminal B, Her2^+^ and TNBC, n=11 in each case) and their functional characteristics were assessed across the subtypes as mentioned in the schematic diagram (Fig. 3A). Distinct gene expression profiles of exhaustion and effector signatures were analyzed through RT-PCR within four BC subtypes. TNBC displayed *eomes^+^ifnγ^low^tox^high^pd1^high^tim3^high^*phenotype, Her2^+^ displayed *eomes^+^ifnγ^moderate^tox^-^pd1^high^tim3^low^*phenotype while Luminal subtypes displayed *eomes^-^ifnγ^high^tox^-^pd1^+^tim3^-^*phenotype (Fig. 3B, 3C).

**Figure 3:**
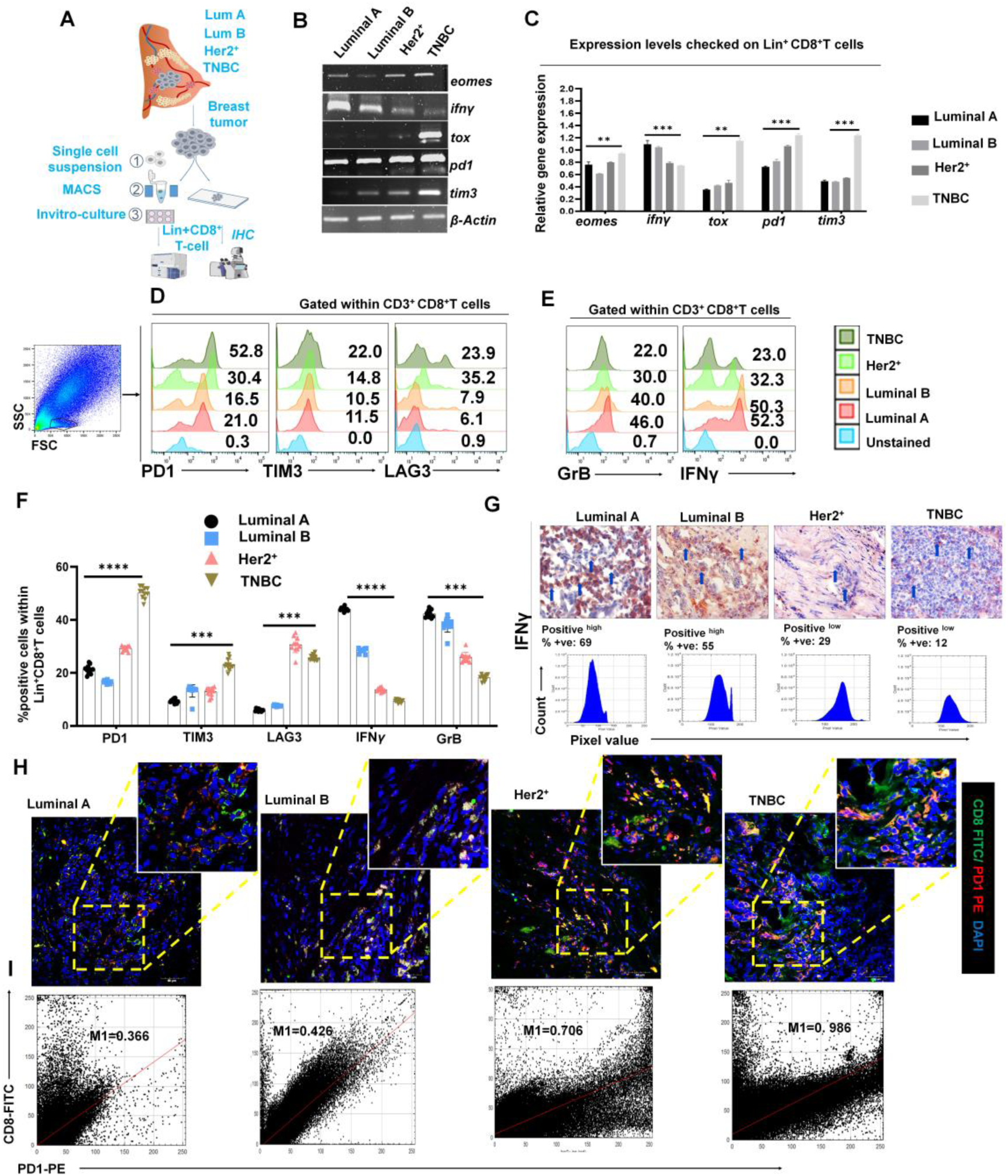
Subtype specific CD8^+^ T-cell infiltration within tumor showcasing distinct-patterns in immune regulation, influencing anti-tumor response. (A) Diagrammatic representation of intratumoral-T cell (Lin^+^CD8^+^) isolation from human BC tissues of the four subtypes. Created in Biorender (See Additional File 5). (B) Identification and genetic-profiling of CD8⁺ T-cell exhaustion and effector pool via RT-PCR from Lin^+^ sorted tumor samples of four subtypes. (C) Gene expression intensities are represented by bar diagrams. Statistical significance estimated from two-way ANOVA followed by Tukey’s multiple comparison (n=3). (D, E) Representative flow-plots for depletion of CD8^+^ T-cell activity and the modulation of the effector cell population generated from Lin^+^ BC samples. (F) Scatter with bar plots represent mean±SD for PD1, TIM3, LAG3, IFNγ, Granzyme B frequency across four subtypes. Statistical significance was drawn by two-way ANOVA followed by Tukey’s multiple comparison test (n=11, in each subtype). (G) Immunohistochemistry was performed in cryopreserved tumor tissue sections (5µm) of IFNγ from four molecular subtypes of BC. IHC profiling was done from Image J IHC profiler and representative histogram plots and log value were calculated by IHC profiler macro. Scoring analysis of the tissue area where the image was captured using a 40X lens and the percentage of the positive and low zones was calculated. Blue arrow lines representing stained zones are provided. Statistical significance inferred from intensity profile generated from Image J. (H) Representative confocal microscopic images at 40X magnification representing co-localization of PD1-PE on CD8-FITC and (within intact tumor sections) of four BC subtypes. (I) Cytofluoromicrographs and Mander’s co-localization co-efficient (M1) for PD1 and co-expression signal are provided below (n=5). (C, F). *\*, p < 0.05; **, p < 0.01; ***, p < 0.001; ****, p < 0.0001*; *ns, nonsignificant* are indicated.

CD8⁺ T cell functional status across BC molecular subtypes was analyzed by flow-cytometry following magnetic isolation of Lin⁺CD8⁺ cells from freshly obtained tumor tissues (Fig. 3D, 3E). Functionally, TNBC exhibited lower IFNγ and Granzyme B with higher PD1 and TIM3. Whereas, heightened expression of IFNγ and Granzyme B were observed in Luminal A and B subtypes. In Her2^+^ tumor, LAG3 expression was slightly elevated compared to TNBC. (Fig. 3D, 3E, 3F).

To identify the functionality of TILs within BC molecular subsets, CD8, PD1 and IFNγ status were evaluated via immunohistochemistry (IHC) and confocal microscopic imaging (Fig. 3G, 3H, 3I). Whole-section IHC analysis revealed that Luminal A and B subtypes are more infiltrated with IFNγ^+^ cells than TNBC and HER2^+^ BCs (Fig. 3G). We next profiled the PD1 expression in CD8⁺ T-cell across intact tumor sections and found that its co-expression was significantly higher in TNBC than other subtypes (Fig. 3H, 3I).

## BCSC mediated CD8⁺ T-cell exhaustion escalates along Luminal to TNBC subtype gradient within TME

Based on prior analyses, BCSC enrichment and intratumoral CD8^+^ T-cell exhaustion markers independently elevated in TNBC, we wanted to further decipher the impact of BCSCs on CD8⁺ T-cells in *in-vitro* co-culture setup. BCSCs and activated CD8^+^ T cells with BCSC: CD8⁺ T-cell in co-culture setup at 1:10 ratio as per schematic diagram (Fig. 4A). BCSC and CD8⁺ T-cells co-culture were performed using two setups (Additional File 1: Supplementary Fig. S2-A); i. Cells were separated through 0.4µm transwell membrane to ensure contact independent communication, ii. Cells were cultured together in same well to confirm contact dependent interaction. The impact of BCSCs on CD8⁺ T-cell were prominent in the contact dependent setup whereas the contact independent study yielded minimal alterations in the functionality of CD8⁺ T-cells.

**Figure 4:**
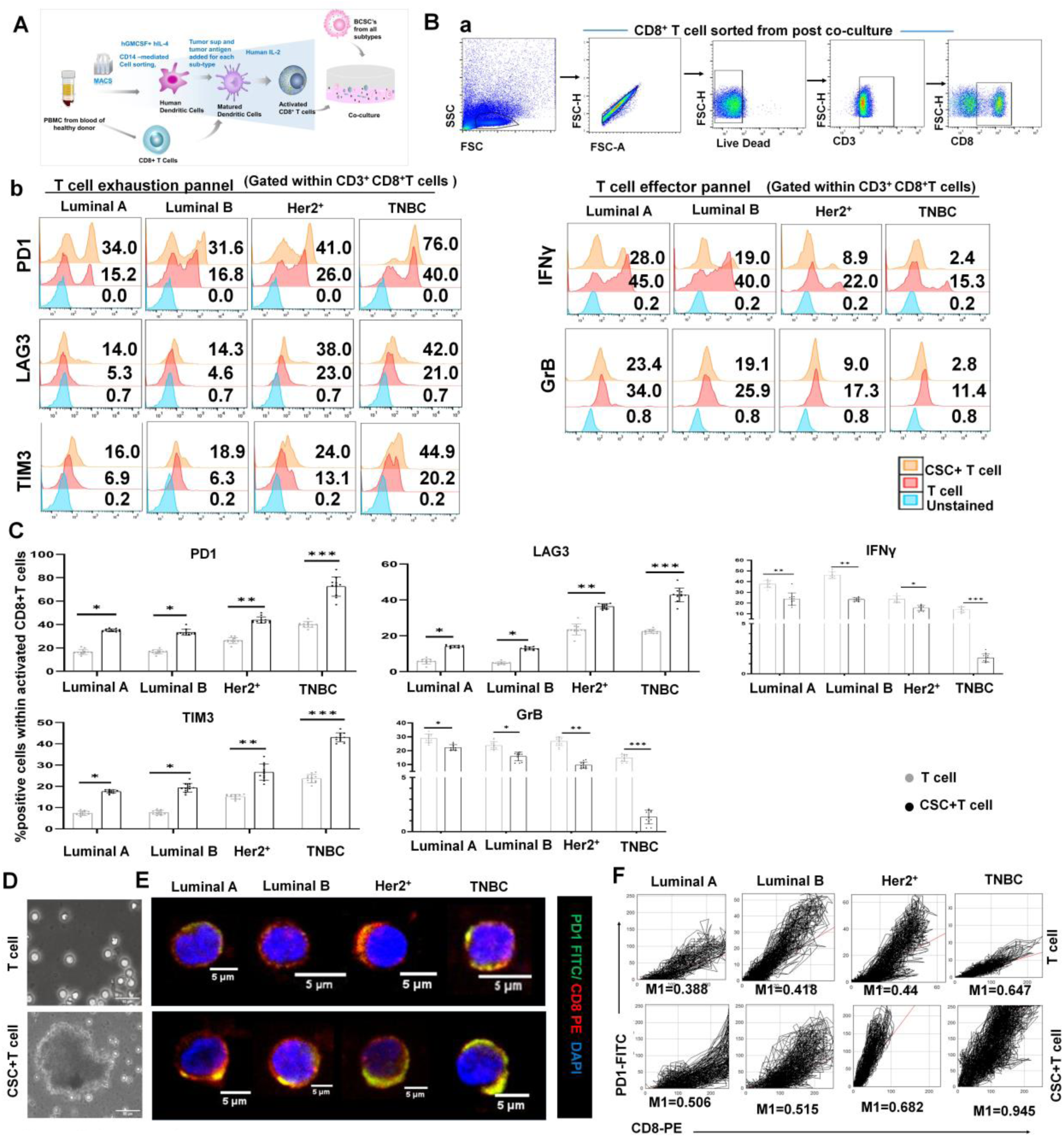
CSC hinders T-cell effector function by promoting their exhaustion to drive aggressiveness. (A) Diagrammatic representation of activated CD8⁺ T-cell generation from human PBMC and their co-culture with BCSCs. Created in Biorender (See Additional File 5). (B) (a) Gating strategy for flow-cytometric analysis. (b) Flow-cytometric histogram plots representing the status of exhaustion (PD1, TIM3, LAG3) and effector (IFNγ, Granzyme B) functions of CD8⁺ T-cells, post co-culture with BCSCs in luminal A&B, Her2^+^ and TNBC subtypes of BC. (C) Scattered with bars displaying relative expression of exhaustion and activation markers in co-culture and untreated group, keeping mean±SD. Statistical significance drawn from two-way ANOVA (n=11) in each case. (D) Representative phase contrast microscopic images at 20X magnification portraying CD8^+^T-cells and physical interaction with BCSCs from patient derived tumor samples (co-cultured with Tu-Ag-pulsed DCs for tumor specific T-cell activation in stem cell enrichment media). (E) Representative confocal microscopic images at 100X magnification representing co-localization of CD8-PE and PD1-FITC (within sorted CD8⁺ T-cells from co-culture of BCSC and CD8⁺ T-cells) of four BC subtypes. (F) Cytofluoromicrographs and Mander’s co-localization co-efficient (M1) for PD1 and co-localization signal are provided below (n=6). (C). *\*, p < 0.05; **, p < 0.01; ***, p < 0.001; ****, p < 0.0001*; *ns, nonsignificant* are indicated.

Next, we explored the functional status of the CD8^+^ T-cells post co-culture via flow-cytometry. Cellular analysis of the CD8⁺ T-cells obtained from each group showed no significant changes in their exhaustion and effector marker status in case of contact-independent co-culture assay (Additional File 1: Supplementary Fig. S2-B). The cytokine status of the T cells also showed no changes across the groups, reaffirming no possible involvement of CSC-secretion mediated exhaustion of CD8⁺ T-cells (Additional File 1: Supplementary Fig. S2-C). Nevertheless, in contact dependent setup, the assessment of functional status of activated T-cells post co-culture revealed significant differences.

Our results suggest that BCSC and T-cell interaction is a physical contact-dependent. From co-culture setup (control T cells and BCSCs co-cultured with CD8^+^ T-cells) of four subtypes, CD8⁺ T-cells were sorted out after 48hrs of culture and analyzed by flow-cytometry followed by confocal microscopy. We assessed the functional status of the activated T-cells post co-culture in detail. The flow-cytometry analysis showed heightened expression of the key exhaustion markers-PD1, LAG3, TIM3 compared to the control T-cells and reduction in expression of IFNγ and granzyme B. Among the subtypes analyzed, CD8⁺ T-cells exposed to TNBC-derived CSCs showed the highest induction of exhaustion marker signatures of effector CD8^+^ T-cells compared to Her2^+^ or Luminal tumor subtypes. Decline in IFNγ and granzyme B was observed in the BCSC conditioned CD8⁺ T-cells across all subtypes. However, the expression significantly fetched down in the TNBC (Fig. 4Ba-b, 4C). These data suggests that in TNBC BCSCs when cultured with CD8⁺ T-cells population, CSCs remain in a more aggravated form and the exhaustion-like phenotype “restrains” the function of CD8⁺ T-cells.

Confocal microscopy demonstrated co-localization of CD8 and PD1 in activated CD8⁺ T-cells, accompanied by a significant increase in PD-1 expression on CD8⁺ T-cells following co-culture. Additionally, PD1 protein level was highest in the TNBC group (Mander’s co-localization co-efficient; M1=0.506 in Luminal A and M1=0.945 in TNBC after co-culture) (Fig. 4D, 4E, 4F). Cumulatively, these data suggested that in TME, BCSCs might enhance the exhaustion of infiltrated CD8⁺ T-cells by ligand-receptor interaction.

## Notch1 orchestrates BCSC expansion and immune evasion via NOTCH1–PD-L1/NANOG axis

Since previous results directed towards contact-dependent effects observed with CD8⁺ T-cell exhaustion for BCSCs, we examined receptor–ligand signaling pathways governing CD8⁺ T-cell programming following direct interaction with BCSCs. Our RT-PCR analysis revealed that *notch1* expression gene was markedly elevated across all the subtypes and highest in TNBC, whereas *notch2, notch3*, *notch4*, *wnt3a, pi3k* and *akt* exhibited no appreciable differences. (Fig. 5A, 5B).

**Figure 5:**
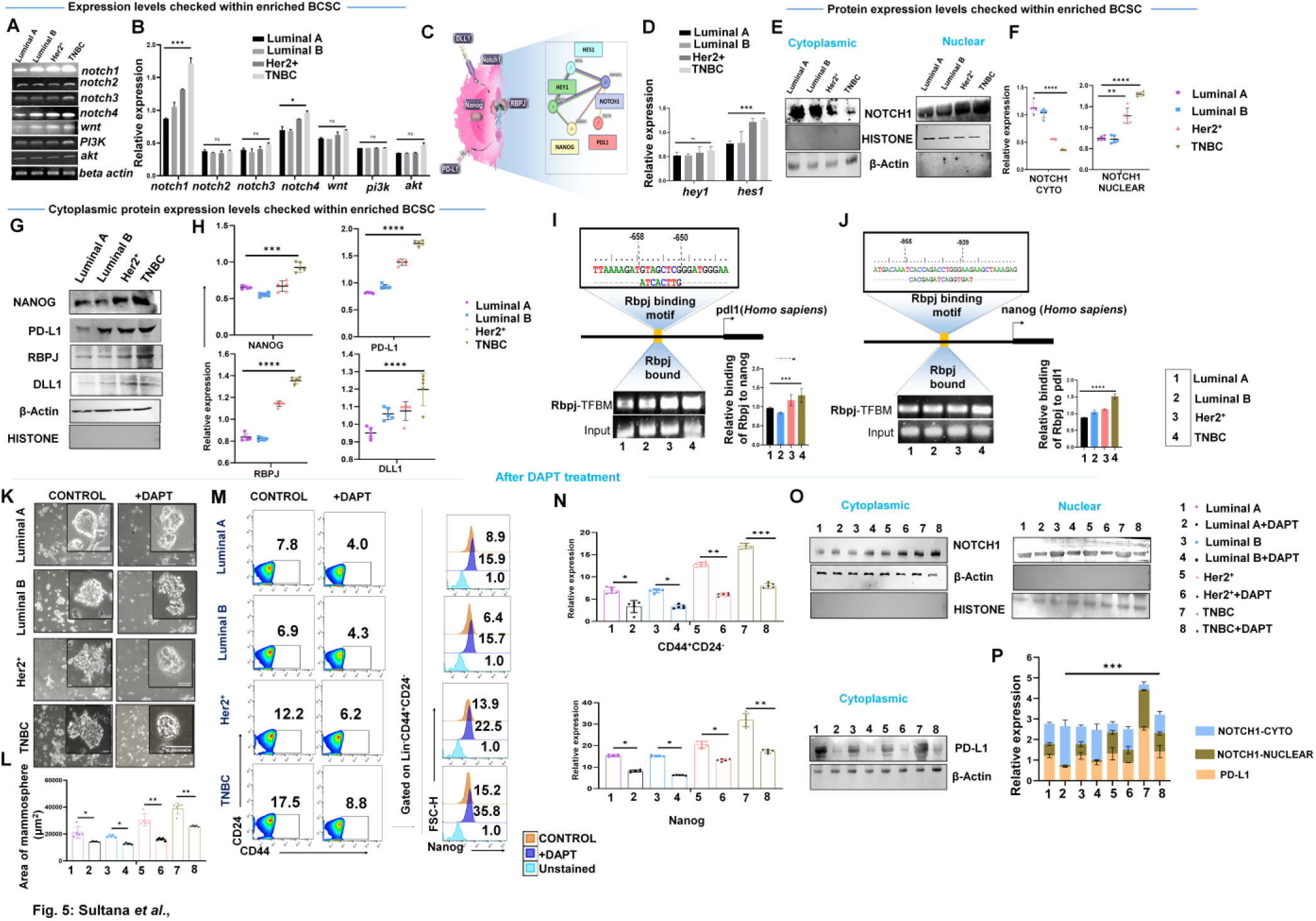
CSC aggressiveness stimulated by Notch1 on BCSC surface, influences through NANOG/PD-L1-PD1 cascade. (A) Analyzing status of CSC cell-surface-receptors regulating BCSC-population. (B) Relative fold-changes in gene expression via bar-graphs (mean± SD); one-way ANOVA followed by Tukey-multiple-comparison test (n=5). (C) General interactions among receptor and probable second messengers of Notch1-association-induced signals in BCSCs, deduced from STRING. (D) Status of *hey1* and *hes1* within BCSC population via RT-PCR. (E) Western blotting to measure NOTCH1 nuclear and cytoplasmic fraction within BCSC (n=5). (F) Scatter plots showcasing protein expression among different subtypes (mean± SD); one-way ANOVA followed by Tukey multiple-comparison test (n = 5). (G) Protein expression within BCSC of all subtypes via Western-blotting. (H) Scatter plots showcasing protein expression among different subtypes (mean± SD); one-way ANOVA followed by Tukey multiple-comparison test (n =5). (I) Pictorial representation of selective binding of Rbpj to their corresponding Transcription-factor-binding-motif (TFBMs) on human *pd-l1* and (J) *nanog* gene due to Notch1 signal. Agarose gel images illustrating the relative amplification of protein-bound TFBMs. Numerical data from different replicates (n=3) are represented. (K) Representative Images of BCSC enriched setups with DAPT treated/untreated cells. (L) Represents bars with scatter plot, for tumorsphere area. (M) Representative pseudocolor-flow-cytometric plots depicting BCSC-frequency and histogram plot depicting Nanog frequency after DAPT treated and untreated cells. (N) Grouped scattered with bars showcasing BCSC-frequency (mean ± SD); one-way ANOVA followed by Tukey multiple-comparison test (n=5). (O) Representative western blots for cytoplasmic and nuclear fractionation DAPT treated and untreated groups. (P) Grouped stacked bars showing protein-expression in different treatment groups (mean ± SD); one-way ANOVA followed by Tukey’s multiple-comparison test (n = 5); (D, F, J, L, N). *\*, p < 0.05; **, p < 0.01; ***, p < 0.001; ****, p < 0.0001*; *ns, nonsignificant* are indicated.

Having ascertained that PD1–mediated immunosuppression underpins the aggressiveness of BCSCs, we further identified a co-ordinated upregulation of *Nanog* and *Notch1* in the BCSC compartment of TNBC. We next sought to predict the possible downstream pathway(s) that may be associated with NOTCH1 and NANOG using the Search Tool for the Retrieval of Interacting Genes/ Proteins (STRING) database. The database showed direct interaction between NOTCH1 and NANOG. Additionally, the database also showed a positive association between PD-L1 and NOTCH1, predicting interplay of NOTCH1 in PD-L1-mediated immune evasion (Fig. 5C).

Assessment of Notch1 downstream molecules revealed that *hes1* was also significantly elevated in TNBC-derived BCSCs, while expression of *hey1* remained unaltered (Fig. 5D). Furthermore, cytoplasmic retention of NOTCH1 in Luminal A, Luminal B and Her2^+^ BC was higher compared to TNBC and notable nuclear localization was observed in all the subtypes, indicating active nuclear translocation suggesting a correlation of Notch1-mediated pathway and degree of aggressiveness of BC (Fig. 5E, 5F). Exploration of NOTCH1 downstream signaling revealed elevated protein level of RBPJ, PD-L1, and NANOG in the aggressive BC subtype TNBC indicating NOTCH1 as a primary regulator in the NOTCH signaling pathway in BCSCs. DLL1, an upstream ligand of Notch1 was also found to be elevated in TNBC subtype further highlighting its aggressive potential (Fig. 5G, 5H). Next, to further investigate possibilities of NOTCH1 mediated transcriptional activation of *nanog* and *pd-l1* genes in BCSC population of four subtypes, we identified transcription factor binding motifs (TFBMs) of RBPJ on the promoter regions of *nanog* and *pd-l1* (*cd274*) using chromatin immunoprecipitation (ChIP) assay. Notch1 effector, RBPJ-bound chromatin complexes were isolated, and specific primers targeting *nanog* and *pd-l1* TFBMs were used for amplification of DNA fragments by RT-PCR. It was revealed that Dll1 induced Notch1 activation with NICD-RBPJ complexes preferentially activates *nanog* transcription and enhances *pd-l1* expression, with highest expression in TNBC CSCs (Fig. 5 I, 5J).

To determine the involvement of Notch1 mediated regulation of Nanog and PD-L1 in human BCSCs, we treated MCF 7 (Luminal A subtype of breast cancer) and MDA-MB-231 (TNBC subtype of breast cancer) with DAPT, a γ-secretase inhibitor (GSI). There mammosphere formation appeared to be enhanced in DAPT-untreated BCSCs, whereas DAPT-treated group showed noticeably smaller mammosphere formation in MCF7 and MDA-MB-231 with prominent changes in MDA-MB-231 cells (Additional File 1: Supplementary Fig. S3-A, B). Additionally, there were 3-fold decrease in BCSC and 2-fold decrease in NANOG expression in presence of DAPT treatment, suggesting NOTCH1-mediated NANOG regulation in human breast cell lines as revealed by flow-cytometric analysis (Additional File 1: Supplementary Fig. S3-C, D). Next, to further validate the above observation in BCSCs of four molecular subtypes, we selectively inhibited Notch1 signaling using DAPT treatment in BCSCs sorted from patient derived tumors. Consistent with earlier observations in cell-line, a significant reduction in mammosphere area and count was observed, particularly in TNBC BCSCs (Fig. 5K, 5L, Additional File 1: Supplementary Fig. S3-E). BCSC percentage inhibition along with reduced NANOG expression was also reflected in all the BCSC subtypes, with the most pronounced effects perceived in TNBC-derived BCSCs (Fig. 5M, 5N). Furthermore, immunolocalization of NOTCH1 via western blot studies revealed a marked decrease in nuclear translocation of Notch1 following DAPT treatment compared to untreated BCSC in TNBC, accompanied by a corresponding increase in cytoplasmic retention of NOTCH1 and downregulation of PD-L1 in DAPT treated group (Fig. 5O, 5P).

Interestingly, CD8⁺ T-cells sorted after co-culture with BCSC also exhibited down regulation of PD1 following DAPT treatment across all the subtypes though the depletion displayed maximum in TNBC (Additional File 1: Supplementary Fig. S4-A, B). Concomitantly, BATF overexpression was observed in TNBC before DAPT treatment, suggesting that CSC enrichment in TNBC may be orchestrated through a PD-L1–PD1 which engages PD1 on CD8⁺ T-cells to activate the PD1–BATF axis. (Additional File 1: Supplementary Fig. S4-C).

## Knockdown of NOTCH1 promotes CD8⁺ T-cell effector functions and suppresses CSC formation in MDAMB-231 cells

Knockdown of Notch1 expression was achieved using specific shRNAs (Fig. 6A). The cloning and transfection process are described in (Additional File 1: Supplementary Fig. S6, S7, S9-S13). Cells transfected with plasmid constructs carrying the different shRNAs, along with the control vector, yielded detectable EGFP expression across all the transfected groups except for the untransfected control (Additional File 1: Supplementary Fig. S7). Initially, we estimated the knockdown efficiency by RT-PCR, in which the Notch1 transcripts were detected in the vector control transfected MDAMB-231 cells, shRNA1, shRNA3 produced negligible reductions and shRNA4 showed high variation among samples. However, shRNA2 consistently showed a ∼50 percent reduction in Notch1 transcripts compared to controls (Fig. 6Ba-b). The downstream genes (*nanog, hes1, pd-l1, and rbpjκ*) in the Notch1 signaling cascade were also analysed by the qRT-PCR analysis. All 4 targets were found to be downregulated in the NOTCH1 knockdown cells based on the level of knockdown of NOTCH1 in the different replicates (Supplementary Fig. S13). This data confirmed the successful knockdown of NOTCH1 in MDA-MB-231 cells. Followed by this, stable NOTCH1 MDAMB-231 knock down (NOTCH1^KD^) cells were generated using lentiviral vectors expressing Notch1 shRNA2. qRT-PCR analysis showed significant knock-down of NOTCH1 expression in stable Notch 1 KD MDAMB-231 cells as compared to wild type cells (Fig. 6Bc-d).

**Figure 6:**
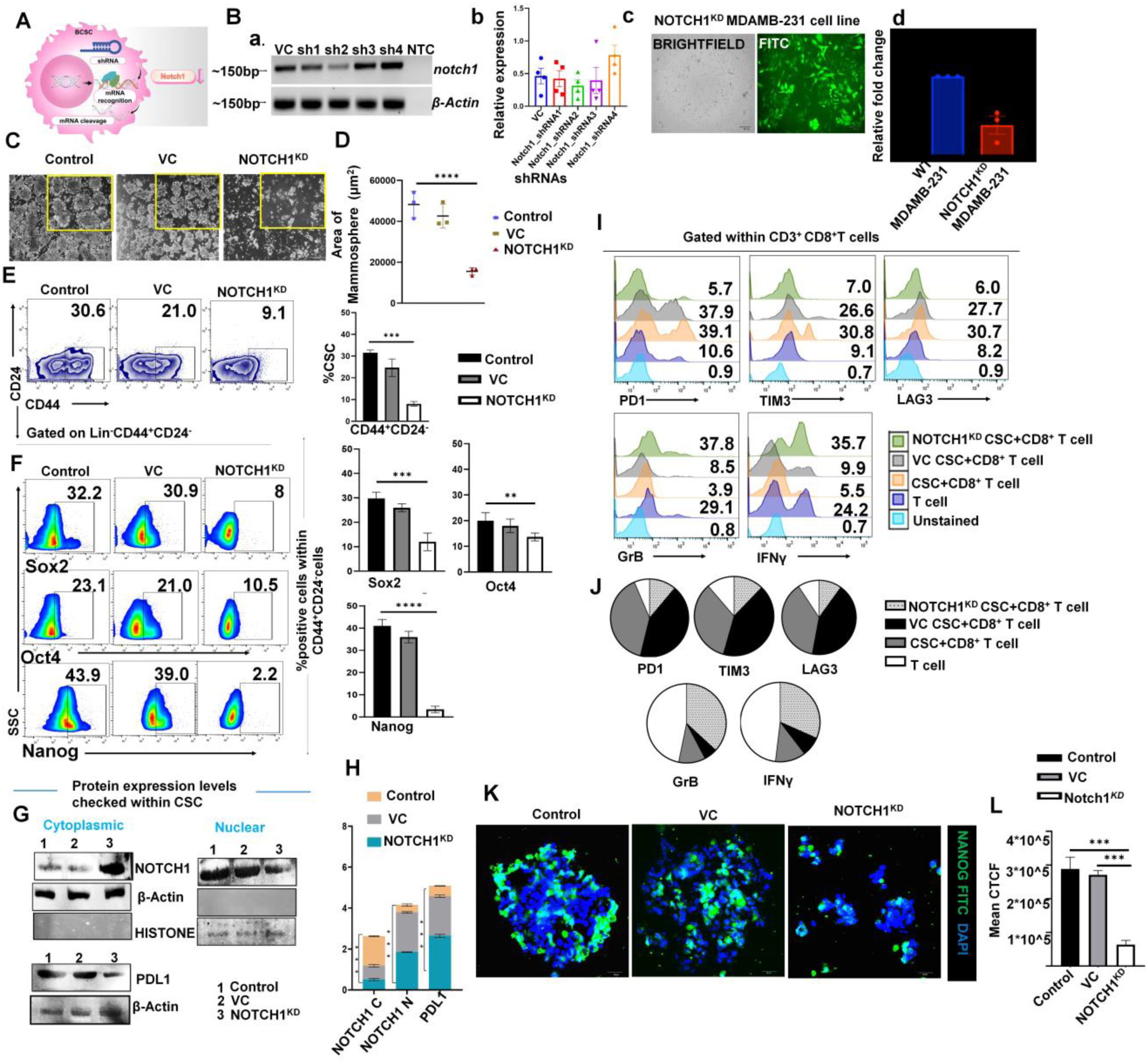
CSC influences CD8^+^ T-cell exhaustion through NOTCH1-NANOG/NOTCH1-PD-L1-PD1 cascade in MDAMB231cells. (A) Schematic illustration of Notch1 knockdown by plasmid-transfection. (B) RT-PCR based quantification of the level of NOTCH1 knockdown in transfected MDAMB-231cells with different NOTCH1-targeting shRNA-constructs, 48 h post-transfection. (a) Agarose gel electrophoresis analysis for the detection of NOTCH1 transcript in the MDAMB-231cells transfected with various shRNAs targeting NOTCH1. VC (vector control); sh1 to sh4 (different-shRNA-expressing-constructs); NTC (no template control). Pictures have been cropped to size for suitable representation. (b) Densitometry analysis of the NOTCH1-expression. The data (mean ± SEM) is indicative of (n=4) independent experiments. (c) fluorescent images of stable NOTCH1^KD^MDAMB-231cells (d) qRT-PCR expression of Notch1 in stable NOTCH1^KD^ MDAMB-231cells as compared to wild type MDAMB-231. The data (mean±SEM) is indicative of (n=3) independent experiments. (C) CSCs plated for tumorsphere assay. Representative images (Control, VC, NOTCH1^KD^) in MDAMB-231cell co-cultures. (D) Scattered graph, mean±SD for tumorsphere-area. Statistical significance is inferred from one-way ANOVA (n=3). (E) Analyzing the status of CSC-fate and (F) stem-cell-regulating-genes within CSC-population via flow-cytometric analysis. Relative fold changes in expression are displayed via bar-graphs (mean±SD); one-way ANOVA (n=3). (G) Western blotting to measure nuclear, cytoplasmic protein suppression by shRNA in MDAMB-231cells (n=3). (H) Statistical significance of stacked bar graphs concluded from one-way ANOVA (n =3) with mean ± SD (right). (I) Flow-cytometric quantifications of proteins within CD8⁺ T-cells of same groups post-culture as half-offset histograms. (J) Representations of all group in each marker by pie-chart. (K) Extent of nuclear localization of NANOG-FITC within mammospheres on silencing Notch(left)(scale bar:20 µm at 100X magnification). (L) Corrected total cell fluorescence intensity (CTCF) as column graph (mean± SD). Statistical alpha value was calculated from unpaired t-test (n=3). *\*, p < 0.05; **, p < 0.01; ***, p < 0.001; ****, p < 0.0001*; *ns, nonsignificant* are indicated.

**Figure 7:**
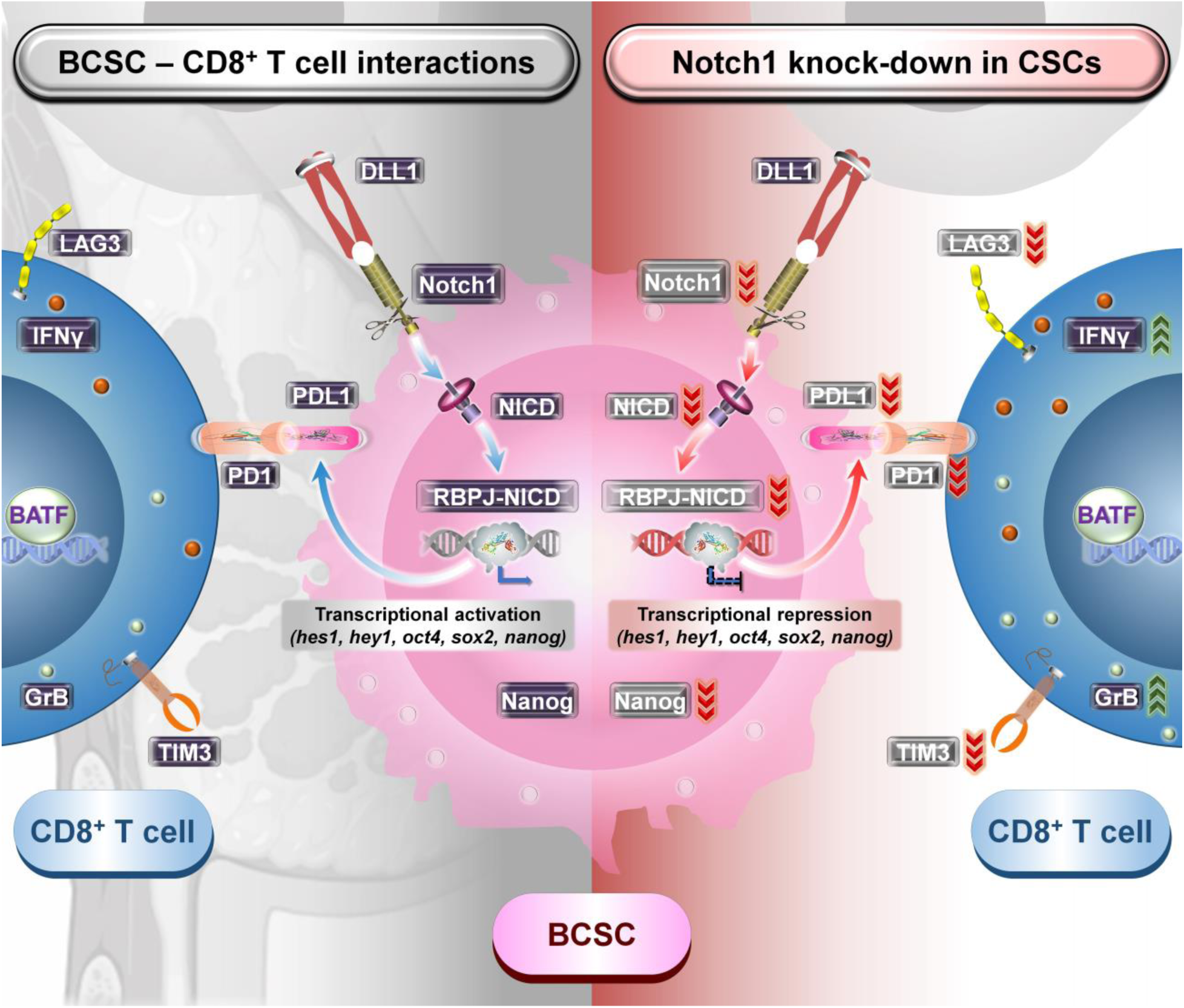
Graphical summary portraying the molecular interactions between BCSCs and CD8^+^ T-cells within the TME alongside the consequences of downregulating Notch1 expressions on BCSCs.

To gain insight into the mechanism of Notch1 upregulation and to determine that whether Notch1 is vital for BCSC survival in MDAMB231 cells, knocked down Notch1 in MDAMB-231 cells were enriched for sorted CSCs along with the unmodified and vector-control MDA-MB-231 cells. Results showed that NOTCH1^KD^MDAMB-231 cells reduced primary mammosphere area formation in shCSC of MDAMB-231 cells by nearly 3-fold compared to control, regardless of culture in enrichment media (Fig. 6C, 6D). Deduction in number of mammosphere was also observed in NOTCH1^KD^MDAMB-231 cells (Supplementary Fig. S14-A). Flow-cytometry was used to check for the expression of CD44^+^CD24^-^ and the intracellular NANOG/OCT4/SOX2 proteins detected by antibody staining. Similar observations were made in NOTCH1^KD^ CSCs, where the CD44⁺/CD24⁻ population exhibited elevated expression of the pluripotency-associated proteins compared with their differentiated counterparts (Fig. 6E). Substantial attenuation of stemness-associated transcription factors was observed, showing 4X reduction in SOX2, 4X downregulation of OCT4, and a pronounced 20X decline in NANOG expression (Fig.s 6F). RT-PCR was used to assess the expression of transcription factors like *sox2, oct4 and nanog* in BCSCs of stable NOTCH1^KD^ MDAMB-231 cells, Vehicle Control (VC) CSC and untreated CSC of MDAMB-231 cells. Expression of stem-cell-related genes confirmed that Notch1^hi^ cells have significantly higher expression of *nanog and sox2*, though changes in *oct4* expression was lower when compared to untreated group (Additional File 1: Supplementary Fig. S14-B).

We next examined the underlying mechanisms of NOTCH1 in regulating immunogenicity. Consistent with our previous observation of reduced PD-L1 following DAPT treatment, NOTCH1^KD^ in MDAMB-231 BCSC cells similarly exhibited markedly decreased PD-L1 expression in a much higher ratio accompanied by reduction in nuclear Notch1 accumulation, relative to untreated and vector-control CSCs (Fig. 6G,6H). Thus, we presumed that NOTCH1 inhibition might augment immunogenicity by downregulating PD-L1/PD1 axis-mediated anticancer immunosuppression upon co-culture of these MDAMB-231 CSCs with CD8⁺ T-cells. Accordingly, co-culture assays were conducted to evaluate this possibility.

We found that co-culture with NOTCH1^KD^MDAMB-231 CSC populations elicited activation phenotype, as evident by heightened IFNγ and Granzyme B with concomitant downregulation of the exhaustion markers PD1, TIM3 and LAG3 in sorted CD8⁺ T-cells post co-culture. Conversely, CD8⁺ T-cells exposed to untreated and vehicle control CSCs showed opposite profile displaying diminished effector functions alongside increased expression of inhibitory receptors (Fig. 6I, 6J).

Next, to further determine whether Notch1 subsidizes to the regulation of Nanog in CSCs of MDAMB-231 in all the three groups (untreated, vehicle control and NOTCH1^KD^ CSCs) was confirmed by immunofluorescence staining. A marked reduction in NANOG expression was observed in the stable NOTCH1^KD^ MDAMB-231 cells suggesting that NOTCH1 may indirectly and positively regulate Nanog expression in CSCs (Fig. 6K, 6L)

## Discussion

Deciphering the molecular features of BCSCs and its corresponding functional dynamics on CD8⁺ T-cells across molecular subtypes of BC remains insufficiently elucidated. BCSC characterization has predominantly been implemented *in-vitro* using established breast cancer cell lines owing to their experimental feasibility (28). While, working with enriched BCSCs offer a more suitable representation of their *in-vivo* tumorigenic and immunomodulatory behavior. Chakravarti et.al. 2023 demonstrated earlier that terminally exhausted cells within breast tumor microenvironment interact with BCSCs and enhances its stemness and metastasis capacity (29). Therefore, we aimed to delineate the distribution of BCSCs and diversified T-cell populations across the molecular subtypes of BC and to observe how BCSCs influence effector T-cell functionality within the TME.

Recent experimental evidences highlight a positive association between BCSCs and CD8⁺ T-cells, coupling stemness-associated signaling with T-cell exhaustion (30). Earlier studies advocates, BCSCs facilitates immune escape by modulating immune checkpoints and cytokine profiles (28,29). However, the mechanistic underpinnings behind BCSC based immunomodulation remains to be explored. Therefore, this study aims to delineate the mechanistic link by which BCSC dampens CD8⁺ T-cell functionality within TME.

Our first analysis majorly focused to study the stem cell proportion and its characterization within molecular subtypes of BC. Flow-cytometry and RT-PCR analysis revealed that TNBC encompasses the highest frequency of BCSCs (CD44⁺CD24⁻) with respect to other molecular BC subtypes. Successively, flow-cytometric analysis of stemness-associated determinants within these subpopulations unveiled an amplified expression of key transcriptional regulators (NANOG), underscoring pronounced stem-like phenotype in TNBC-derived BCSCs compared to other less aggressive molecular subtypes.

This result aligns with the evidence supporting high Nanog expression in TNBC samples during immunohistochemistry analysis (31–34). Similar observation was noted upon further enrichment of BCSCs with stem cell enrichment media. These findings reinforce prior reports stating that certain oncogenic signal pathway may be involved in modulation of BCSC properties through Nanog within cancer (35). More the passages the higher is the tumorigenicity in TNBC derived BCSCs (i.e., mammosphere-forming efficiency, both in sphere count and area). Corresponding, flow-cytometric profiling and confocal imaging further escalated expression of stemness signatures. NANOG expression heightened in TNBC, tailed by HER2-enriched, Luminal A, and Luminal B cohorts as passages advanced, revealing a passage-dependent reinforcement of the pluripotent circuitry. BCSCs develop several strategies within TME to evade immune surveillance, by reducing infiltration of cytotoxic T-lymphocytes and increasing employment of immunosuppressive cells, as reported by other studies (36–38). BCSCs and their immune microenvironment are interwoven entities and CD8⁺ T-cells play pivotal role in tumor restriction (39). Therefore, next we investigated the CD8⁺ T-cell characteristics, within BC and their effector functions using flow-cytometry. Our findings revealed that TNBC subtypes characterized by high expression of PD1, TIM3, LAG3 (inhibitory receptors) in all the subtypes. PD1, LAG3, TIM3 within CD8⁺ T-cells were negatively correlated with the low prevalence of immune effector signatures such as IFNγ and Granzyme B. This observation was detected across molecular subtypes, suggesting an exceedingly high immunogenic TME that supports the rationale for targeting this immune checkpoint blockade (40–42). Immunohistochemistry of PD1 and IFNγ also supports the similar observation across molecular subtypes of BC. Furthermore, to evaluate precise impact of BCSC’s on CD8^+^ T-cell exhaustion, we co-cultured cancer stem cells with activated CD8^+^ T-cells. Previously, it has been demonstrated that upregulation of stemness regulated genes like OCT4, SOX2, NANOG within BCSC results in escape from specific anti-tumor immunity provided by CD8^+^ T-cells (43,44). In line with their report, here we have found that in post co-culture a marked upregulation of exhaustion markers (PD1, TIM3, LAG3) were observed, followed by a concomitant reduction in CD8⁺ T-cells effector molecules (IFNγ and Granzyme B) using flow-cytometry. This dysregulation in immune profile was most elevated in BCSC populations derived from TNBC, implicating BCSCs as significant modulator of CD8⁺T cells dysfunction and immune evasion in aggressive BC subtypes.

Understanding critical insights into how BCSC impacts immune evasion, is an important strategy to counteract BCSC survival and recurrence. Collectively, our results demonstrated that TNBC-enriched BCSCs harbors distinct NANOG –driven stemness reservoir and upon BCSC culture with CD8⁺ T-cells, they concomitantly elicited proportion of PD1 on CD8⁺ T-cells. Hence, we intended to find a link between them via STRING analysis. This network indicated pivotal role of NOTCH signaling axis, connecting it as a potential molecular conduit synchronizing stemness growth with immune-checkpoint activation. So, next we checked expression of several NOTCH associated molecules and involvement of NOTCH1 was observed which matches well with previous evidences which identifies NOTCH1 as a powerful target predominantly for BCSCs (45–48). HES1, an important downstream target gene of NOTCH was found to be predominantly upregulated in TNBC. Beside this, RBPJ and DLL1, key arbitrators of NOTCH transcriptional activation and ligand interaction respectively, were observed to be at the highest levels in TNBC BCSCs compared to the other BC subtypes. Moreover, upregulation of NOTCH1-DLL1 juxtracrine signaling leads to the expansion of BCSCs (49). Additionally, emerging evidences indicate that the oscillatory pattern of NOTCH1 and PD-L1 are linked. NOTCH1 contributes to immune-evasive phenotypes through upregulation of PD-L1 and thereby can support BCSC survival and immune escape (50–52).Mechanistic studies confirmed the RBPJ (key transcriptional effector of the NOTCH1 signaling pathway) binding to the promoters of PD-L1 and NANOG, establishing that NOTCH1 mediates BCSC upregulation through NANOG induction while concurrently promoting immune suppression via the PD-L1/PD1 axis. Inhibition of Notch1 signaling by gamma-secretase inhibitor such as DAPT downsizes tumor-sphere formation and normalizes downstream immune suppression. In agreement with our previous result, NOTCH1 was markedly overexpressed in TNBC samples. Several shRNAs were constructed to make NOTCH1^KD^MDAMB-231 for confirmation of involvement of NOTCH1 mediated immunosuppression. Knockdown of NOTCH1 using shRNA2 construct in the MDAMB-231 cell line, a representative model of TNBC, gave rise to knockdown of NOTCH1 transcripts, suggesting it was the most effective among the four designs. The level of knockdown achieved was sufficient to induce preliminary changes in downstream genes such as Nanog, Hes1, RBPJ, and PD-L1, indicating biological relevance. These genes are critical for stemness, transcriptional regulation, and immune evasion, respectively. Reduction in BCSC population, lowered the expression of stemness-associated transcription factors, this decline was prominent in expression of Nanog. This downregulation further aligned with reduced PD-L1 expression, subsidizing to the attenuation of T-cell exhaustion through modulation of PD1 signaling. Collectively, these findings propose that therapeutic inhibition of NOTCH1 may effectively target CSC-mediated tumor progression in TNBC by concurrently impairing stemness preservation and immune evasion mechanisms. PD1 induces the expression of the transcription factor BATF that sustains PD1 expression and enforces T-cell exhaustion.

Altogether, this study provides mechanistic insights into how NOTCH1 signaling pathway, differentially modulate BCSC population to drive tumor aggressiveness in different subtypes of BC. TNBC exhibits the most pronounced activation of these regulatory pathways, consistent with its highly aggressive nature. Our findings suggest that the ultimate therapeutic efficacy depends on both the degree of stemness induction by NANOG and other transcription factors and the exhausted state of infiltrating immune cells like PD1, TIM3, and LAG3.

This study demonstrates the subtype-specific distribution of BCSCs among molecular subtypes of BC and also deciphers their immune crosstalk with CD8⁺ T-cells. By mapping how aggressive TNBC-derived BCSCs orchestrate T-cell dysfunction, this work provides mechanistic insight into stemness-immunity coupling within an Asian cohort. However, different geographic contexts may yield distinct immuno-biological signatures. Moreover, our observations suggest that neutralizing or targeting DLL1/PD-L1-NOTCH1/NANOG on BCSCs within the TME has a possible benefit to counteract-immune-checkpoint inhibition.

## Conclusions

Immunotherapy as a single approach often fails to generate durable responses in solid tumors due to intrinsic immune resistance within TME. Notably, inhibiting Notch1 in BCSCs increases cytotoxic T-cell infiltration and sensitizes tumors to PD1/PD-L1–directed therapies. These findings support the rationale that dual targeting of Notch1-driven stemness and checkpoint pathways may provide a promising combinatorial treatment to enhance immunotherapeutic responses in aggressive BC subtypes.

## Supplementary Information

### Additional file 1

Supplementary Figures S1-S15. Fig.S1. Supporting information of Figure 2 stating that BCSC increases with passages. Fig.S2. Supporting information of Figure 4. BCSC-CD8^+^ T-cell co-culture direct towards a physical contact-dependent CD8^+^ T-cell exhaustion mediated by BCSC. Fig.S3. Supporting information of Figure 5 showing the DAPT treated downregulation in MCF-7 and MDAMB-231 cells. Fig.S4. Continued supporting data for Figure 5 showing PD1 expression within CD8^+^ T-cell sorted from BCSC-CD8^+^ T-cell co-culture with or without DAPT treatment. Fig.S5. Schematic representation of the shRNA cloning strategy in pRNAT-CMV3.1-Neo vector. Fig.S6. Cloning of shRNA Oligonucleotides into pRNAT-CMV3.1-NeoVector. Fig.S7. Fluorescence microscopy imaging of EGFP expression in MDA-MB-231 cells upon transfection with different Notch1 shRNA constructs. Fig.S8. Schematic representation of generation of Notch1 shRNA2 expression construct (pCMV-EGFP miR-E Notch1 shRNA2) containing pCMV-EGFP miR-E backbone. Fig.S9. Cloning of Notch1 shRNA2 into pCMV-EGFP miR-E Vector. Fig.S10. Transfection of MDA-MB-231 cells with pCMV-EGFP miR-E Notch1 shRNA2 clones. Fig.S11. Schematic representation of the CMV-EGFP miR-E Notch1 shRNA2 expression cassette in pLJM1-EGFP vector. Fig.S12. Sub-cloning of CMV-EGFP miR-E shRNA2 expression in pLJM1-EGFP vector. Fig.S13. Quantitative Real time PCR to analyze the level of expression of the downstream genes of Notch1. Fig.S14. Supporting information for Figure 6 showing downregulation of mammosphere count and stemness associated genes.

### Additional file 2

Supplementary Tables T1-T5. Table T1. Reagent and antibody details. Table T2. Details of human patient sample. Table T3. Sequences and primer list of genes. Table T4. List of shRNA sequences used in the study. Table T5. List of primer and their seqence used in the study for knockdown experiments.

### Additional file 5

Merged pdf of the Confirmation of Publication and Licensing Rights of biorender are provided.

## Declaration

## Supporting information

Biorender license

## Abbreviations

AEC: 3-amino-9-ethylcarbazole
AKT: Protein kinase B
ANOVA: Analysis of variance
BATF: Basic leucine zipper ATF-like transcription factor
BC: Breast cancer
BCSC: Breast cancer stem cells
CaBr: Breast cancer
CD24: Cluster of differentiation 24
CD44: Cluster of differentiation 44
CD47: Cluster of differentiation 47
CD274: Cluster of differentiation 274 (PD-L1)
ChIP: Chromatin immunoprecipitation
CSC: Cancer stem cells
DAPT: N-[N-(3,5-difluorophenacetyl-L-alanyl)]-S-phenylglycine t-butyl ester
DC: Dendritic cell
DLL1: Delta-like canonical Notch ligand 1
EGFP: Enhanced green fluorescent protein
ELISA: Enzyme-linked immunosorbent assay
EOMES: Eomesodermin
ER: Estrogen receptor
FITC: Fluorescein isothiocyanate
GrB: Granzyme B
GSI: γ-secretase inhibitor
HES1: Hairy and enhancer of split 1
HER2: Human epidermal growth factor receptor 2
HEY1: Hairy/enhancer-of-split related with YRPW motif protein 1
HIF-1α: Hypoxia-inducible factor 1 alpha
IFN-γ: Interferon gamma
IHC: Immunohistochemistry
IL2: Interleukin 2
IL10: Interleukin 10
IL12: Interleukin 12
IL13: Interleukin 13
KD: Knockdown
KLRG1: Killer cell lectin-like receptor G1
LAG3: Lymphocyte activation gene 3
LIN: Lineage
LUMA: Luminal A
LUMB: Luminal B
MACS: Magnetic-activated cell sorting
MDA-MB-231: MD Anderson metastatic breast cancer cell line
MCF-7: Michigan Cancer Foundation-7 cell line
MDSC: Myeloid-derived suppressor cell
MHC-I: Major histocompatibility complex class I
mRNA: Messenger ribonucleic acid
NACT: Neoadjuvant chemotherapy
NF-κB: Nuclear factor kappa B
NICD: Notch intracellular domain
NTC: No-template control
OCT4: Octamer-binding transcription factor 4
PBMC: Peripheral blood mononuclear cell
PD-1: Programmed cell death protein 1
PD-L1: Programmed death ligand 1
PD-L2: Programmed death ligand 2
PE: Phycoerythrin
PI3K: Phosphoinositide 3-kinase
PR: Progesterone receptor
qPCR: Quantitative polymerase chain reaction
qRT-PCR: Quantitative reverse transcription polymerase chain reaction
RBPJ: Recombination signal binding protein for immunoglobulin kappa J region
RT-PCR: Reverse transcription polymerase chain reaction
SD: Standard deviation
shRNA: Short hairpin ribonucleic acid
SHP-2: Src homology 2 domain-containing protein tyrosine phosphatase 2
SOX2: SRY-box transcription factor 2
STAT3: Signal transducer and activator of transcription 3
STRING: Search tool for the retrieval of interacting genes/proteins
TAM: Tumor-associated macrophage
TFBM: Transcription factor binding motifs
TGF-β: Transforming growth factor beta
TIL: Tumor-infiltrating lymphocytes
TIM-3: T-cell immunoglobulin and mucin domain 3
TME: Tumor microenvironment
TNBC: Triple-negative breast cancer
TNF-α: Tumor necrosis factor alpha
TOX: Thymocyte selection-associated high mobility group box
TREG: Regulatory T cell
TCR: T-cell receptor
TSS: Transcription start site
TU-AG: Tumor antigen
VC: Vehicle control
VEGF: Vascular endothelial growth factor
WNT: Wingless-related integration site

## Acknowledgements

Besides institutional aid, this work was supported by Indian Council of Medical Research (ICMR) New Delhi to J.S. (File No: 3/2/2/63/2022-NCD-III), University Grants Commission (UGC), New Delhi to P.R.C. (Ref. No: 191620061322), Council of Scientific and Industrial Research (CSIR), New Delhi to S.Be. (File No: 112-1831-6176/2K23/1) and Indian Council of Medical Research (ICMR), New Delhi to S.B. (File No: 5/13/21/2022/NCD-III). Funding only supports fellowship to scholars and cost of experimental reagents. The authors cordially thank Director CNCI, Kolkata, India for providing institutional facilities. Special thanks to Director of BRIC-National Institute of Animal Biotechnology, Hyderabad and National Institute for Pharmaceutical Education & Research (NIPER), Mohali for their collaboration with the project. The authors thank Rupankar Ghosh for his persistent support in flow-cytometric analysis. The authors also thank all respective laboratory members for their constant support. The authors declare that they have not use AI-generated work in the manuscript

## Consent for Publication

All authors have given their consent to participate

## Data availability

No data sets were analyzed during the study. All data analyzed during this study will be available from the corresponding author on reasonable request.

## Authors’ contributions

S. Ba., A. B., and J.S. designed and directed all experiments. S. Ba., A. B., M.C. and J.S. oversaw all experiments, wrote and reviewed the manuscript. N.Ga provided the facility for knockdown experiments. A.G., P.R.C, P.D. assisted with cell culture. S.Be., J.D. assisted with histological experiments. G.U. conducted the knockdown and performed the associated analyses. S.Be., J.D., N.G assisted with schematic figures and graphical abstracts. M.C., S.D., A.S., N.G., supported preliminary experiments. M.S., M.R. assisted in performing knockdown. A.Sa. gifted antibodies. M.C., P.R.C., P.D., S.Be, A.G., S.D., A.S., N.G., A.Sa, helped in editing the manuscript. N.A. supervised clinical aspects of the study. J.S., S.Ba, A. B. and R.B prepared figures analyzed and edited data. Overall proposal was comprehended by S.Ba.

## Ethics Approval

All human experiments were approved by Institutional Ethical Committee of Chittaranjan National Cancer Institute, Kolkata, India. Approval number (Approval no: CNCI-IEC-SB-20) dated 25.04.2022, under the topic titled as “Understanding the influence of transcription factors on cancer stem cells from different molecular subtypes of breast cancer in remodeling of tumor immune landscape: Therapeutic intervention by 2DG and NLGP”.

## Consent to Participate

Human breast tumor and blood samples were collected following written consent from patients and institutional ethical committee. Approval number (Approval no: CNCI-IEC-SB-20).

## References

1. Horr C, Buechler SA. Breast Cancer Consensus Subtypes: A system for subtyping breast cancer tumors based on gene expression. npj Breast Cancer. 2021 Oct 12;7(1):136.

2. Romaniuk-Drapa³a A, Totoñ E, Taube M, Idzik M, Rubiœ B, Lisiak N. Breast Cancer Stem Cells and Tumor Heterogeneity: Characteristics and Therapeutic Strategies. Cancers (Basel). 2024 Jul 7;16(13):2481.

3. Zhong M, Ren X, Xia W, Qian Y, Sun K, Wu J. The role of adjuvant endocrine treatment in ER+, PR−, HER2− early breast cancer: a retrospective study of real-world data. Sci Rep. 2024 Nov 2;14(1):26377.

4. Ghosh A, Chaubal R, Das C, Parab P, Das S, Maitra A, et al. Genomic hallmarks of endocrine therapy resistance in ER/PR+HER2-breast tumours. Commun Biol. 2025 Feb 10;8(1):207.

5. Tommasi C, Airò G, Pratticò F, Testi I, Corianò M, Pellegrino B, et al. Hormone Receptor-Positive/HER2-Positive Breast Cancer: Hormone Therapy and Anti-HER2 Treatment: An Update on Treatment Strategies. J Clin Med. 2024 Mar 24;13(7):1873.

6. Chakraborty S, Basak U, Mukherjee S, Mukherjee S, Das T. Cancer Stem Cells Decide the Fate of Cancer Immunotherapy by Remodeling Tumor Microenvironment. Cancer Control. 2025 Sep 24;32.

7. Guo Q, Zhou Y, Xie T, Yuan Y, Li H, Shi W, et al. Tumor microenvironment of cancer stem cells: Perspectives on cancer stem cell targeting. Genes Dis. 2024 May;11(3):101043.

8. Gao Q, Zhan Y, Sun L, Zhu W. Cancer Stem Cells and the Tumor Microenvironment in Tumor Drug Resistance. Stem Cell Rev Reports. 2023 Oct 21;19(7):2141–54.

9. Gómez-Aleza C, Nguyen B, Yoldi G, Ciscar M, Barranco A, Hernández-Jiménez E, et al. Inhibition of RANK signaling in breast cancer induces an anti-tumor immune response orchestrated by CD8+ T cells. Nat Commun. 2020 Dec 10;11(1):6335.

10. Zhang J, Liu B, Lyu M, Duan Y. Cutting the root: the next generation of T cells engagers against cancer stem cells to overcome drug resistance in triple-negative breast cancer. Cancer Biol Med. 2023 Mar 24;20(3):169–73.

11. Syrnioti A, Petousis S, Newman LA, Margioula-Siarkou C, Papamitsou T, Dinas K, et al. Triple Negative Breast Cancer: Molecular Subtype-Specific Immune Landscapes with Therapeutic Implications. Cancers (Basel). 2024 May 31;16(11):2094.

12. Lin Y, Song Y, Zhang Y, Li X, Kan L, Han S. New insights on anti-tumor immunity of CD8+ T cells: cancer stem cells, tumor immune microenvironment and immunotherapy. J Transl Med. 2025 Mar 17;23(1):341.

13. Zhong H, Zhou Z, Wang H, Wang R, Shen K, Huang R, et al. The Biological Roles and Clinical Applications of the PI3K/AKT Pathway in Targeted Therapy Resistance in HER2-Positive Breast Cancer: A Comprehensive Review. Int J Mol Sci. 2024 Dec 13;25(24):13376.

14. Xu T, Zhang H, Yang BB, Qadir J, Yuan H, Ye T. Tumor-infiltrating immune cells state-implications for various breast cancer subtypes. Front Immunol. 2025 May 14;16.

15. Hou Y, Nitta H, Wei L, Banks PM, Lustberg M, Wesolowski R, et al. PD-L1 expression and CD8-positive T cells are associated with favorable survival in HER2-positive invasive breast cancer. Breast J. 2018 Nov;24(6):911–9.

16. Xie H, Xi X, Lei T, Liu H, Xia Z. CD8+ T cell exhaustion in the tumor microenvironment of breast cancer. Front Immunol. 2024 Dec 9;15.

17. Rouzbahani E, Majidpoor J, Najafi S, Mortezaee K. Cancer stem cells in immunoregulation and bypassing anti-checkpoint therapy. Biomed Pharmacother. 2022 Dec;156:113906.

18. Bergdorf KN, Phifer CJ, Bechard ME, Lee MA, McDonald OG, Lee E, et al. Immunofluorescent staining of cancer spheroids and fine-needle aspiration-derived organoids. STAR Protoc. 2021 Jun;2(2):100578.

19. Schneider CA, Rasband WS, Eliceiri KW. NIH Image to ImageJ: 25 years of image analysis. Nat Methods. 2012 Jul 28;9(7):671–5.

20. Varghese F, Bukhari AB, Malhotra R, De A. IHC Profiler: An Open Source Plugin for the Quantitative Evaluation and Automated Scoring of Immunohistochemistry Images of Human Tissue Samples. Aziz SA, editor. PLoS One. 2014 May 6;9(5):e96801.

21. Nair S, Archer GE, Tedder TF. Isolation and Generation of Human Dendritic Cells. Curr Protoc Immunol. 2012 Nov;99(1).

22. Ganguly N, Das T, Bhuniya A, Guha I, Chakravarti M, Dhar S, et al. Neem leaf glycoprotein binding to Dectin-1 receptors on dendritic cell induces type-1 immunity through CARD9 mediated intracellular signal to NFκB. Cell Commun Signal. 2024 Apr 23;22(1):237.

23. Dhar S, Chakravarti M, Ganguly N, Saha A, Dasgupta S, Bera S, et al. High monocytic MDSC signature predicts multi-drug resistance and cancer relapse in non-Hodgkin lymphoma patients treated with R-CHOP. Front Immunol. 2024 Jan 18;14.

24. Fellmann C, Hoffmann T, Sridhar V, Hopfgartner B, Muhar M, Roth M, et al. An Optimized microRNA Backbone for Effective Single-Copy RNAi. Cell Rep. 2013 Dec;5(6):1704–13.

25. Ulgekar G, Kaur D, Ganesan V, Sen Sharma S, Ganguli N, Majumdar SS. Anhydride chemistry-based hexanoylation of polyethylenimine increases transfection efficiency and expression of tagged DNA for therapeutic proteins in cultured cells. Biotechnol Bioeng. 2022 Nov 6;119(11):3275–83.

26. Ali K, Nabeel M, Mohsin F, Iqtedar M, Islam M, Rasool MF, et al. Recent developments in targeting breast cancer stem cells (BCSCs): a descriptive review of therapeutic strategies and emerging therapies. Med Oncol. 2024 Apr 9;41(5):112.

27. Baldasici O, Soritau O, Roman A, Lisencu C, Visan S, Maja L, et al. The transcriptional landscape of cancer stem-like cell functionality in breast cancer. J Transl Med. 2024 Jun 3;22(1):530.

28. Xie F, Zhou X, Su P, Li H, Tu Y, Du J, et al. Breast cancer cell-derived extracellular vesicles promote CD8+ T cell exhaustion via TGF-β type II receptor signaling. Nat Commun. 2022 Aug 1;13(1):4461.

29. Chakravarti M, Dhar S, Bera S, Sinha A, Roy K, Sarkar A, et al. Terminally Exhausted CD8+ T Cells Resistant to PD-1 Blockade Promote Generation and Maintenance of Aggressive Cancer Stem Cells. Cancer Res. 2023 Jun 2;83(11):1815–33.

30. Hu H, Zou M, Hu H, Hu Z, Jiang L, Escobar D, et al. A breast cancer classification and immune landscape analysis based on cancer stem-cell-related risk panel. npj Precis Oncol. 2023 Dec 8;7(1):130.

31. Tian Y, Zhang P, Mou Y, Yang W, Zhang J, Li Q, et al. Silencing Notch4 promotes tumorigenesis and inhibits metastasis of triple-negative breast cancer via Nanog and Cdc42. Cell Death Discov. 2023 May 6;9(1):148.

32. Pierantozzi E, Gava B, Manini I, Roviello F, Marotta G, Chiavarelli M, et al. Pluripotency Regulators in Human Mesenchymal Stem Cells: Expression of NANOG But Not of OCT-4 and SOX-2. Stem Cells Dev. 2011 May;20(5):915–23.

33. Placzek S, Vanzan L, Deluz C, Suter DM. Orchestration of pluripotent stem cell genome reactivation during mitotic exit. Cell Rep. 2025 Apr;44(4):115486.

34. Nagata T, Shimada Y, Sekine S, Moriyama M, Hashimoto I, Matsui K, et al. KLF4 and NANOG are prognostic biomarkers for triple-negative breast cancer. Breast Cancer. 2017 Mar 14;24(2):326–35.

35. Vasefifar P, Motafakkerazad R, Maleki LA, Najafi S, Ghrobaninezhad F, Najafzadeh B, et al. Nanog, as a key cancer stem cell marker in tumor progression. Gene. 2022 Jun;827:146448.

36. Peng D, Tanikawa T, Li W, Zhao L, Vatan L, Szeliga W, et al. Myeloid-Derived Suppressor Cells Endow Stem-like Qualities to Breast Cancer Cells through IL6/STAT3 and NO/NOTCH Cross-talk Signaling. Cancer Res. 2016 Jun 1;76(11):3156–65.

37. Wu B, Shi X, Jiang M, Liu H. Cross-talk between cancer stem cells and immune cells: potential therapeutic targets in the tumor immune microenvironment. Mol Cancer. 2023 Feb 21;22(1):38.

38. Mukherjee S, Chakraborty S, Basak U, Pati S, Dutta A, Dutta S, et al. Breast cancer stem cells generate immune-suppressive T regulatory cells by secreting TGFβ to evade immune-elimination. Discov Oncol. 2023 Dec 1;14(1):220.

39. Guha A, Goswami KK, Sultana J, Ganguly N, Choudhury PR, Chakravarti M, et al. Cancer stem cell–immune cell crosstalk in breast tumor microenvironment: a determinant of therapeutic facet. Front Immunol. 2023 Nov 27;14.

40. Yakubovich E, Cook DP, Rodriguez GM, Vanderhyden BC. Mesenchymal ovarian cancer cells promote CD8+ T cell exhaustion through the LGALS3-LAG3 axis. npj Syst Biol Appl. 2023 Dec 12;9(1):61.

41. Sadeghi Rad H, Monkman J, Warkiani ME, Ladwa R, O’Byrne K, Rezaei N, et al. Understanding the tumor microenvironment for effective immunotherapy. Med Res Rev. 2021 May 4;41(3):1474–98.

42. Wei J, Li W, Zhang P, Guo F, Liu M. Current trends in sensitizing immune checkpoint inhibitors for cancer treatment. Mol Cancer. 2024 Dec 26;23(1):279.

43. Feng D, Pu D, Ren J, Liu M, Zhang Z, Liu Z, et al. CD8+ T-cell exhaustion: Impediment to triple-negative breast cancer (TNBC) immunotherapy. Biochim Biophys Acta – Rev Cancer. 2024 Nov;1879(6):189193.

44. Noh KH, Kim BW, Song KH, Cho H, Lee YH, Kim JH, et al. Nanog signaling in cancer promotes stem-like phenotype and immune evasion. J Clin Invest. 2012 Nov 1;122(11):4077–93.

45. Uribe-Etxebarria V, Pineda JR, García-Gallastegi P, Agliano A, Unda F, Ibarretxe G. Notch and Wnt Signaling Modulation to Enhance DPSC Stemness and Therapeutic Potential. Int J Mol Sci. 2023 Apr 17;24(8):7389.

46. Zhang Y, Xu W, Guo H, Zhang Y, He Y, Lee SH, et al. NOTCH1 Signaling Regulates Self-Renewal and Platinum Chemoresistance of Cancer Stem–like Cells in Human Non–Small Cell Lung Cancer. Cancer Res. 2017 Jun 1;77(11):3082–91.

47. Wang LL, Wan XY, Liu CQ, Zheng FM. NDR1 increases NOTCH1 signaling activity by impairing Fbw7 mediated NICD degradation to enhance breast cancer stem cell properties. Mol Med. 2022 Dec 4;28(1):49.

48. Zhang C, Xu S, Yin C, Hu S, Liu P. The role of the mTOR pathway in breast cancer stem cells (BCSCs): mechanisms and therapeutic potentials. Stem Cell Res Ther. 2025 Mar 29;16(1):156.

49. Shah PA, Huang C, Li Q, Kazi SA, Byers LA, Wang J, et al. NOTCH1 Signaling in Head and Neck Squamous Cell Carcinoma. Cells. 2020 Dec 12;9(12):2677.

50. Montagner A, Arleo A, Suzzi F, D’Assoro AB, Piscaglia F, Gramantieri L, et al. Notch Signaling and PD-1/PD-L1 Interaction in Hepatocellular Carcinoma: Potentialities of Combined Therapies. Biomolecules. 2024 Dec 11;14(12):1581.

51. Cao Y, Liu B, Cai L, Li Y, Huang Y, Zhou Y, et al. G9a promotes immune suppression by targeting the Fbxw7/Notch pathway in glioma stem cells. CNS Neurosci Ther. 2023 Sep 27;29(9):2508–21.

52. Wang M, Yu F, Zhang Y, Li P. Novel insights into Notch signaling in tumor immunity: potential targets for cancer immunotherapy. Front Immunol. 2024 Feb 20;15.

