## Supplementary Figures S1-S14 for "Breast cancer stem cells mediated CD8^+^ T cell exhaustion among different molecular subtypes of breast cancer regulated via NOTCH1/RBPJ/PD-L1 axis"

**Dr. Saptak Banerjee**

Senior Scientific Officer

Department of Immunoregulation and Immunodiagnostics

Chittaranjan National Cancer Institute

37, S.P. Mukherjee Road

Kolkata – 700026, India

Contact: +91-9717615367

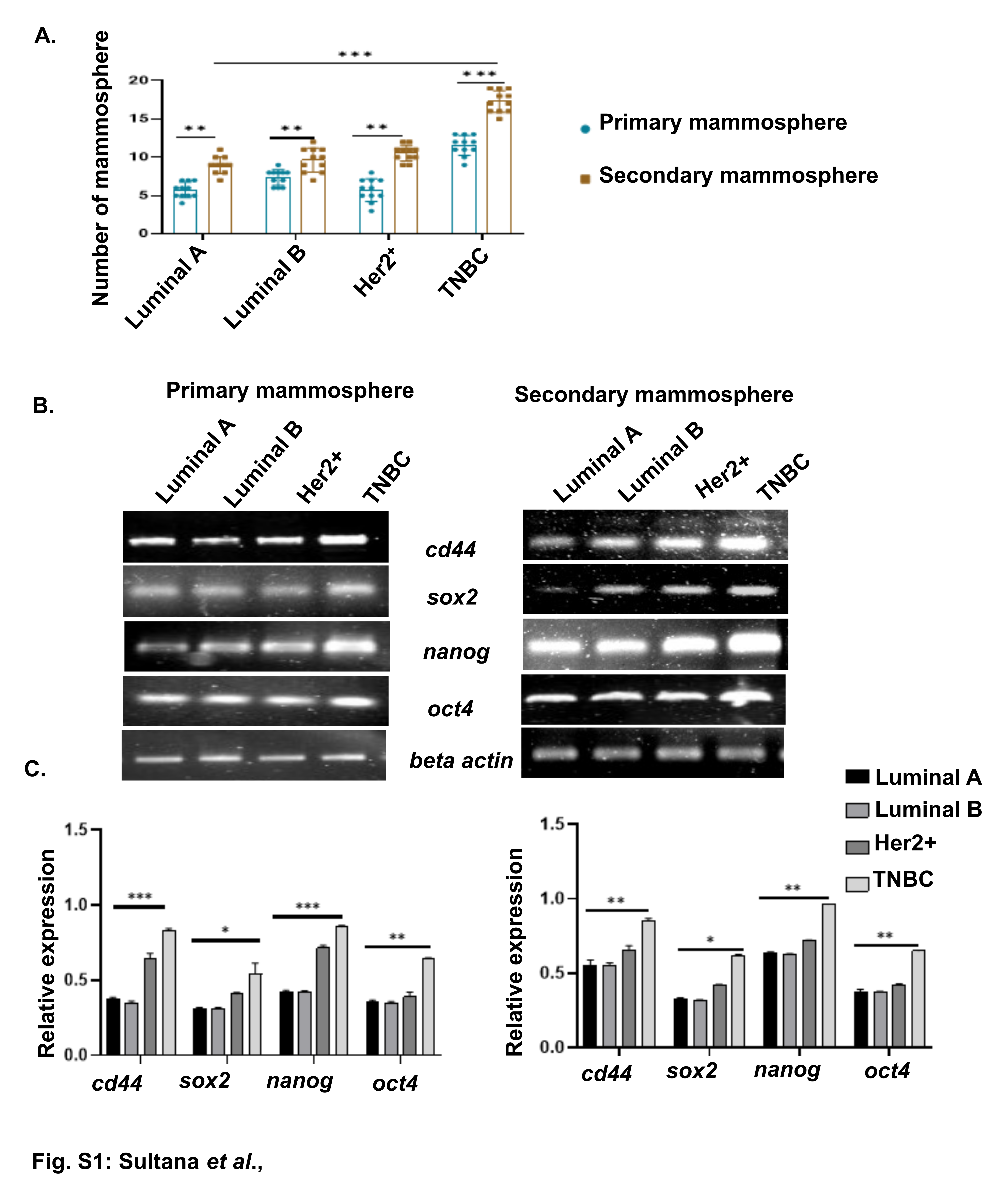

**Supplementary Figure S1**: (A) In bar-graph, mean±SD for tumorsphere count is indicated. Statistical significance is inferred from two-way ANOVA followed by Tukey’s multiple comparison test (n=11). (B) Relative fold changes in *cd44, sox2, nanog* and *oct 4* expression within CSC cultured in stem cell enrichment media. Relative fold changes in the bars (mean± SD), (C) Statistical significance drawn from two-way ANOVA followed by Tukey’s multiple comparison test (n=5). (A, C) **, p < 0.05; **, p < 0.01; ***, p < 0.001; ****, p < 0.0001*; *ns, nonsignificant* are indicated.

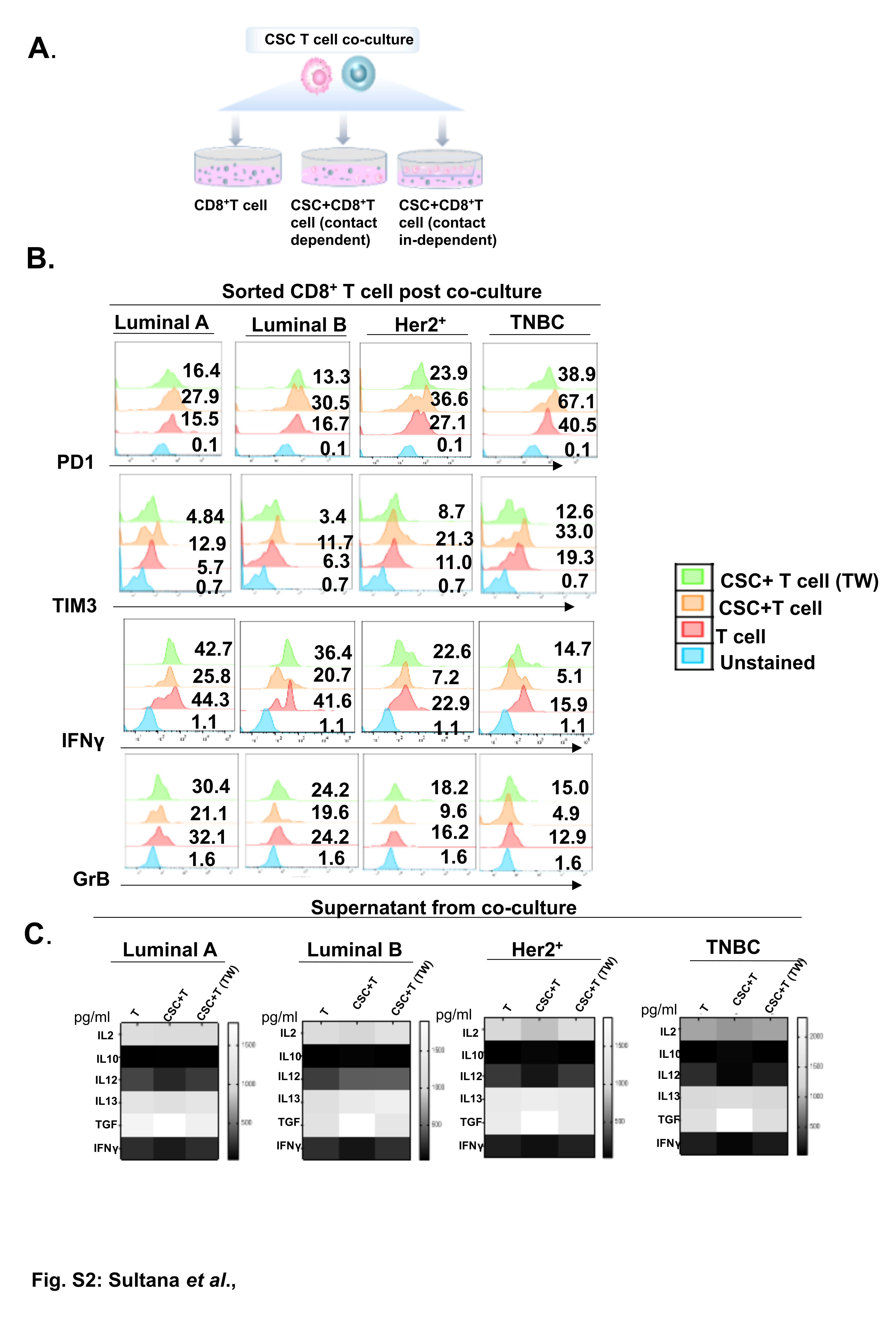

**Supplementary Figure S2:** (A) Illustrative representation for probable scenarios of BCSC mediated CD8^+^T cell exhaustion. (B) Representative histogram plots representing changes in CD8^+^T cell exhaustion and effector functions in co-culture setups of four subtypes of BC, with and without transwell membrane. CD8T cell with no treatment kept as control. (C) Evaluation of possible alteration in cytokines due to co-culture between BCSC (Luminal A, Luminal B, Her2^+^ and TNBC) and CD8^+^T cells by ELISA. Co-culture supernatants were assessed against no-treatment CD8^+^T cells. Each box in heatmap represents mean±SD for pg/ml cytokine secretion (n=3) where dark color indicates low production and light color indicates high production.

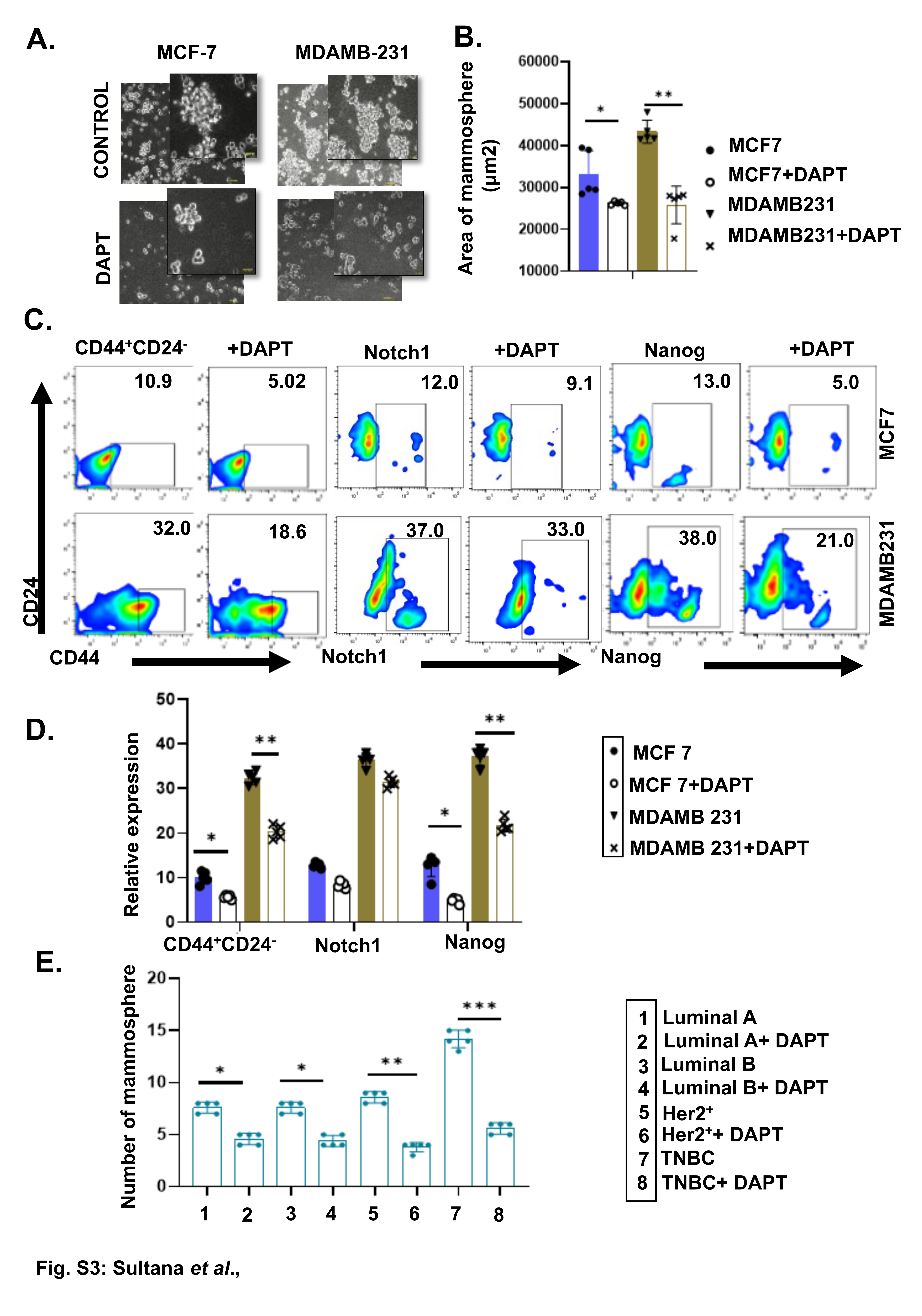

**Supplementary Figure S3:** (A) Representative images of CSC of MCF7 and MDAMB-231 cell line with and without DAPT treated cultures are provided. (B) Scattered with bar graphs representing the area of mammosphere across all experimental groups (mean ± SD); one-way ANOVA followed by Tukey multiple-comparison test (n =5). (C) Representative pseudocolor flow-cytometric plots for, BCSC, Notch1 and Nanog for all experimental groups. (D) Scattered with bar graphs showcasing percentage positive cell frequency in various treated and untreated groups (mean ± SD); one-way ANOVA followed by Tukey multiple-comparison test (n=5). (E) In bar-graph, mean±SD for tumorsphere count is indicated with and without DAPT treatment in BCSC of four subtypes. Statistical significance is inferred from two-way ANOVA followed by Tukey’s multiple comparison test (n=5). (B, D, E) **, p < 0.05; **, p < 0.01; ***, p < 0.001; ****, p < 0.0001*; *ns, nonsignificant* are indicated.

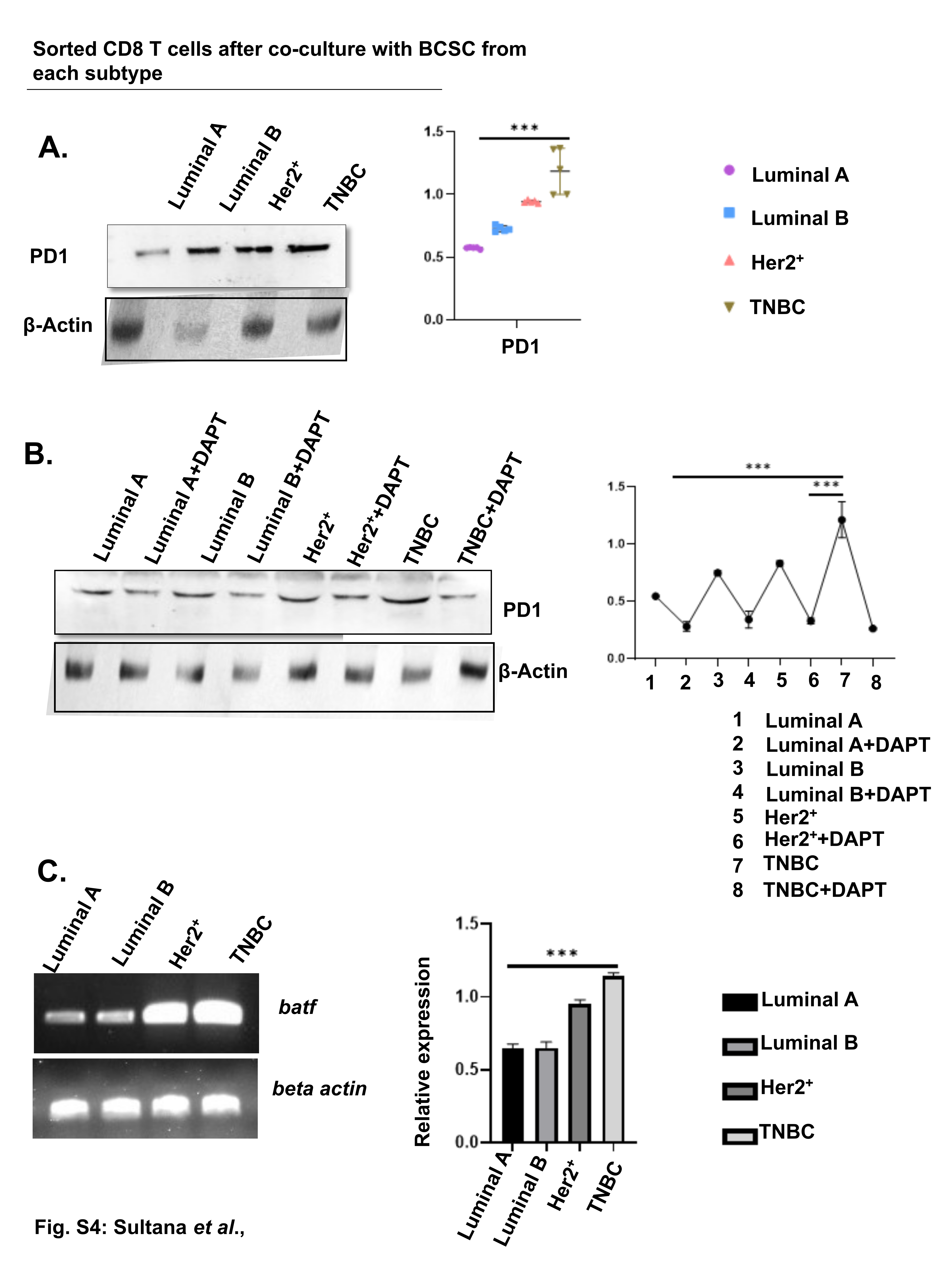

**Supplementary Figure S4:** (A) Representative western blots for protein fractions of PD-1 in the 4 subtypes. Grouped scatter plot showcasing variant protein expression. Grouped scatter plot showcasing variant protein expression of in the corresponding treatment sets (mean±SD); Evaluation measured by one-way ANOVA analysis followed by Tukey’s multiple-comparison-test (n=5). (B) PD-1 protein expression after DAPT (pharmacological inhibitor of Notch) treatment in four subtypes. Symbols with connecting line showcasing variant protein expression of in the corresponding treatment group (mean±SD); Analysis done by one-way ANOVA analysis followed by Tukey’s multiple-comparison-test (n=5). (C) Analyzing the trancription factor *batf* (induced by PD-1) status within CD8^+^ T cells via qPCR. Relative fold changes in gene expression across all groups are displayed via bar-graphs (mean± SD); one-way ANOVA analysis followed by Tukey’s multiple-comparison-test (n=5). Full-length uncropped-blot presented in Additional-file4. (A, B, C) **, p < 0.05; **, p < 0.01; ***, p < 0.001; ****, p < 0.0001*; *ns, nonsignificant* are indicated.

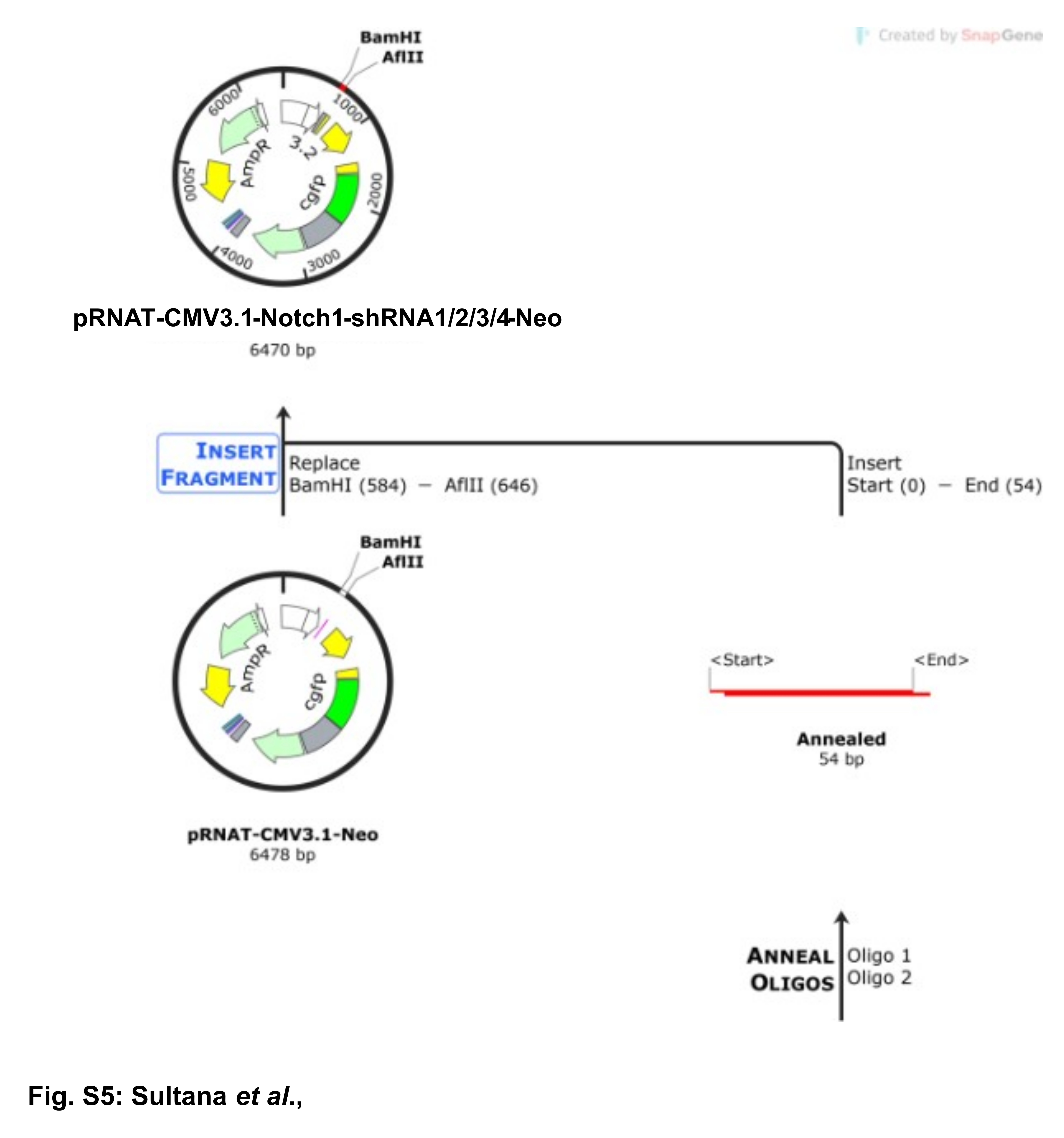

**Supplementary Figure S5:** Schematic representation of the shRNA cloning strategy in pRNAT-CMV3.1-Neo vector.

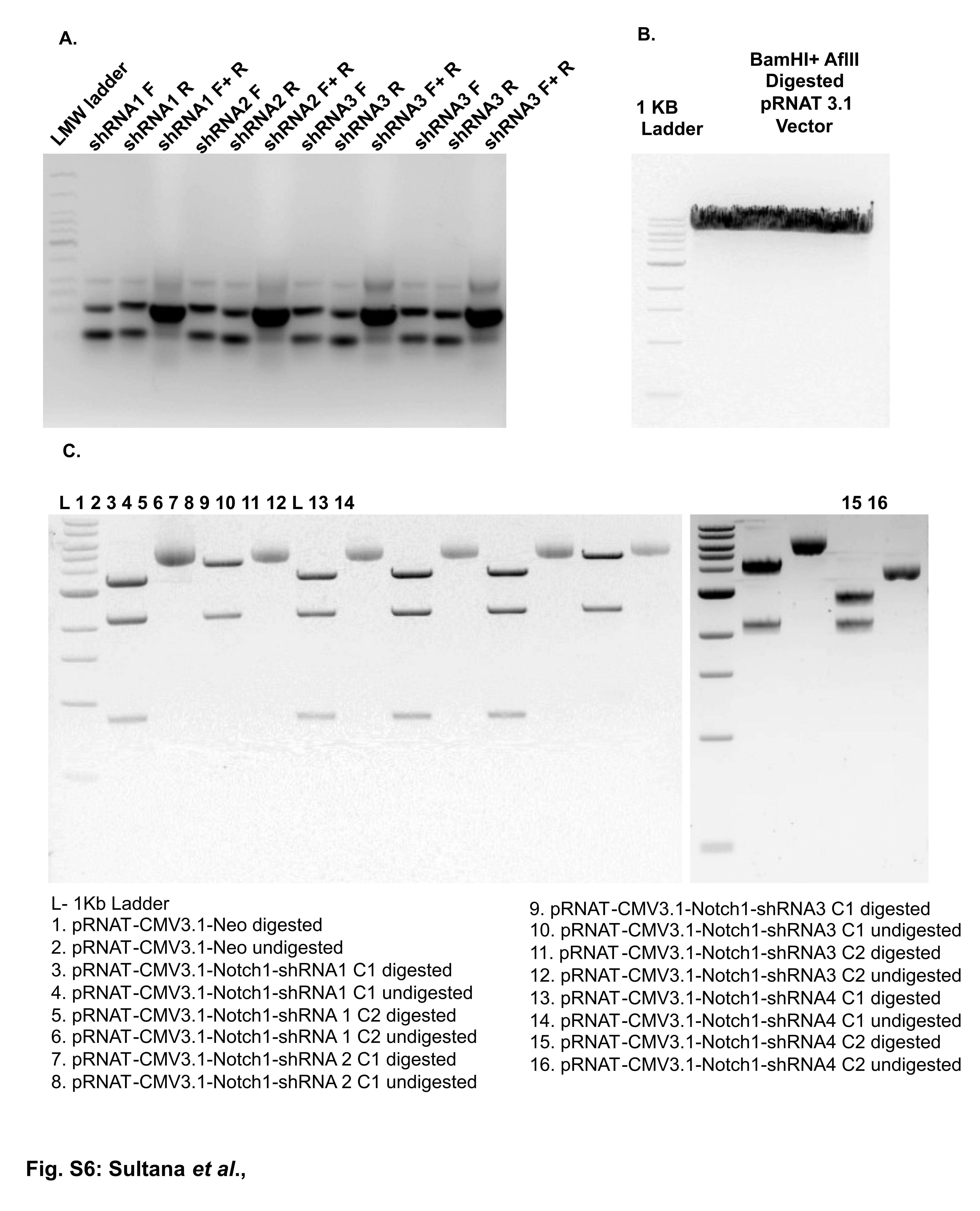

**Supplementary Figure S6:** Cloning of shRNA Oligonucleotides into pRNAT-CMV3.1-NeoVector.
(A) Annealing of shRNA oligonucleotides for insertion.
(B) Restriction digestion of the pRNAT-CMV3.1-Neo vector using BamHI and AflII for ligation.
(C) Restriction mapping of potential clones using SalI and EcoRV to confirm successful insertion.

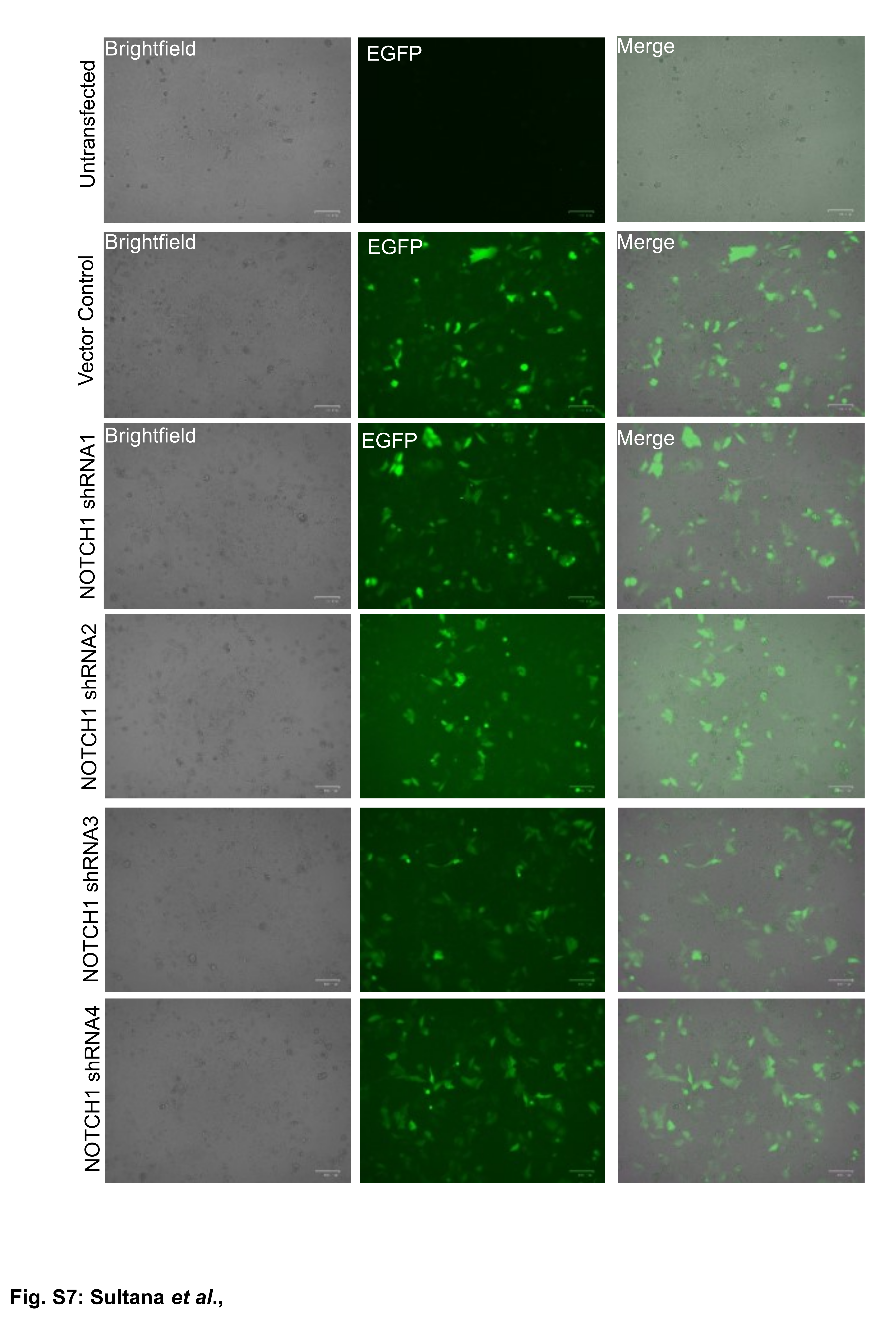

**Supplementary Figure S7:** Fluorescence microscopy imaging of EGFP expressed in MDA-MB-231 cells upon transfection with different Notch1 shRNA constructs, 48 hours post transfection.

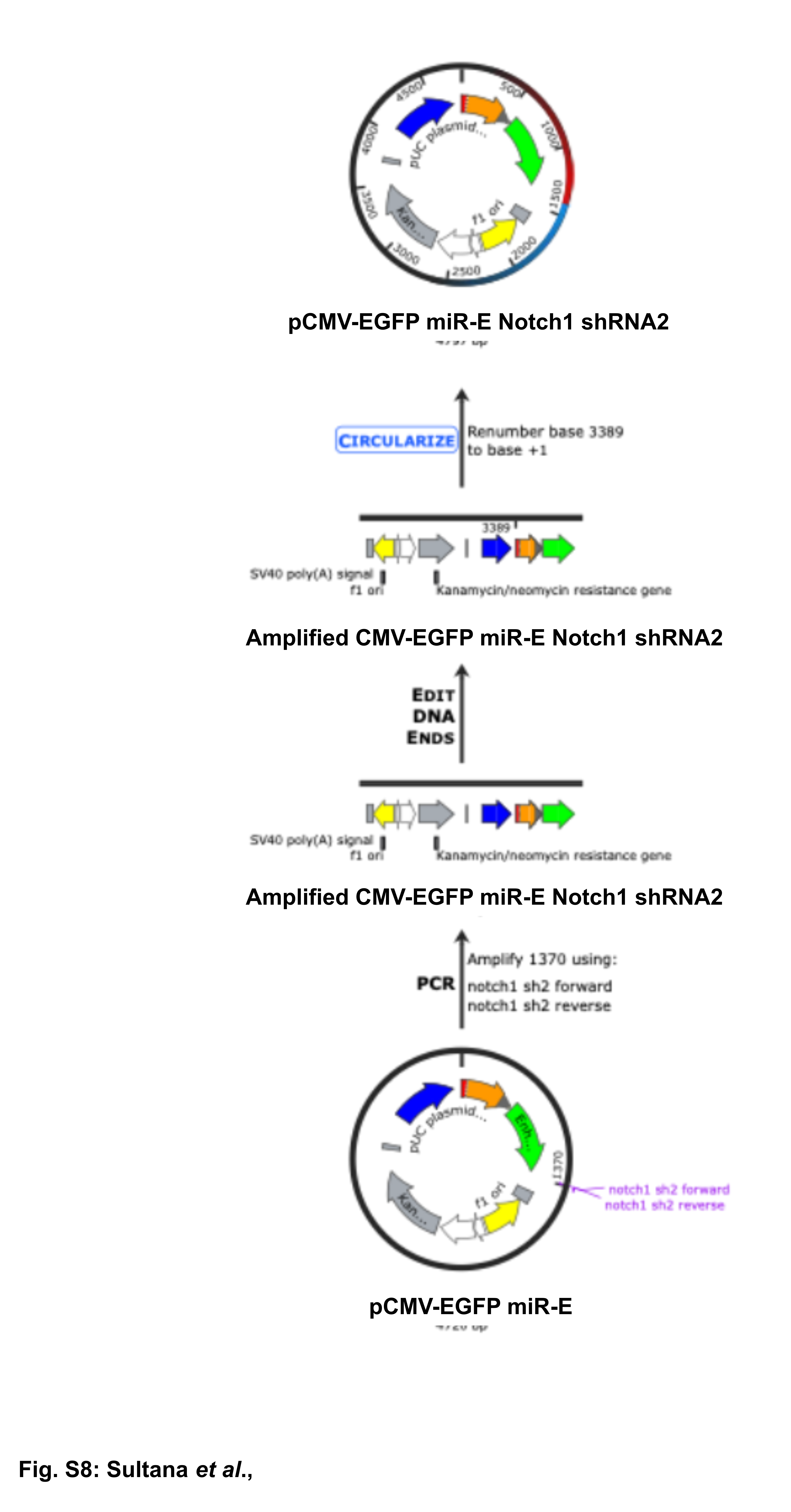

**Supplementary Figure S8:** Schematic representation of generation of Notch1 shRNA2 expression construct (pCMV-EGFP miR-E Notch1 shRNA2) containing pCMV-EGFP miR-E backbone.

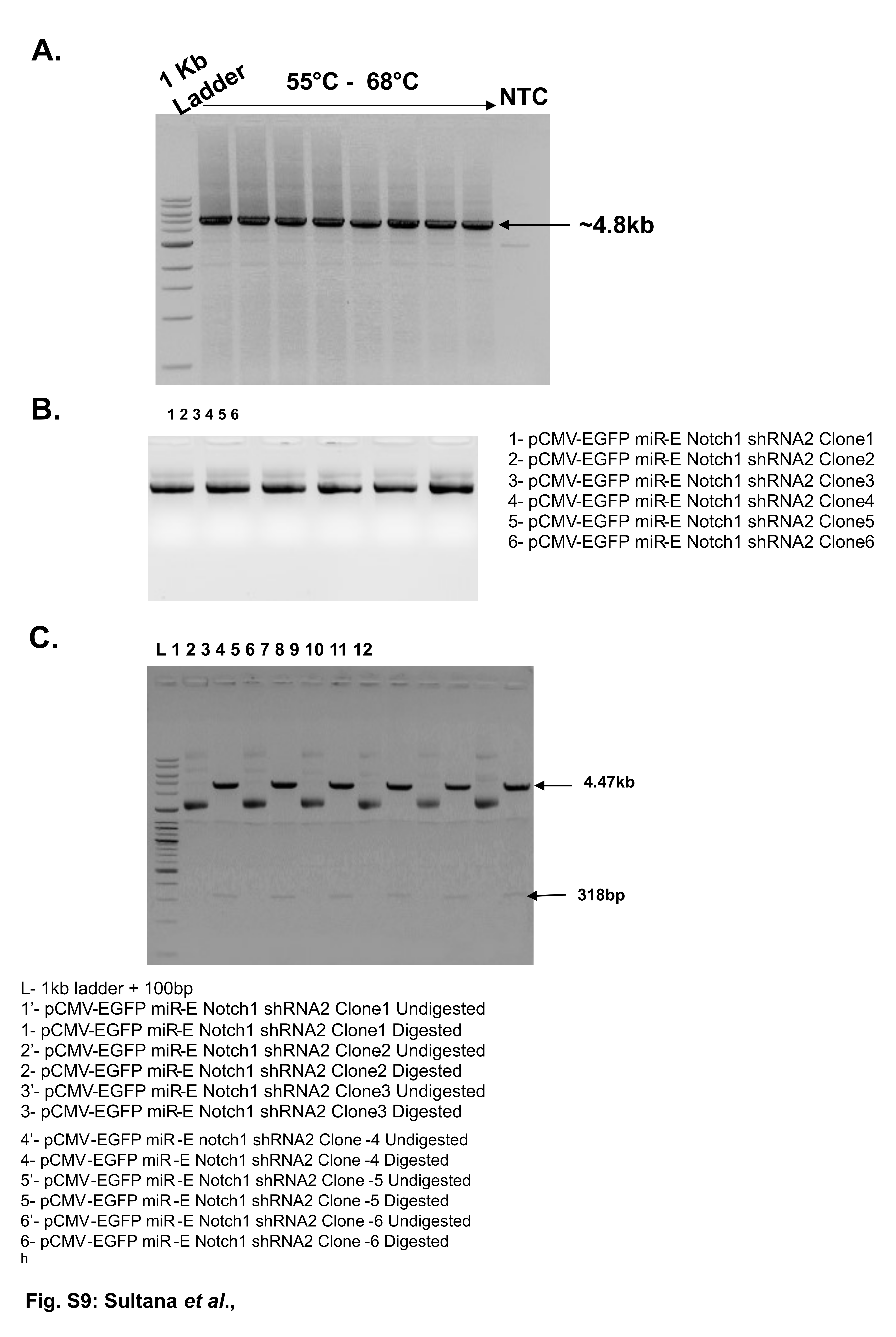

**Supplementary Figure S9:** Cloning of Notch1 shRNA2 into pCMV-EGFP miR-E Vector.
(A) Amplification of pCMV-EGFP miR-E Notch1 shRNA2.
(B) Isolation of plasmids from potential positive colonies.
(C) Restriction mapping of recombinant clones to confirm successful insertion.

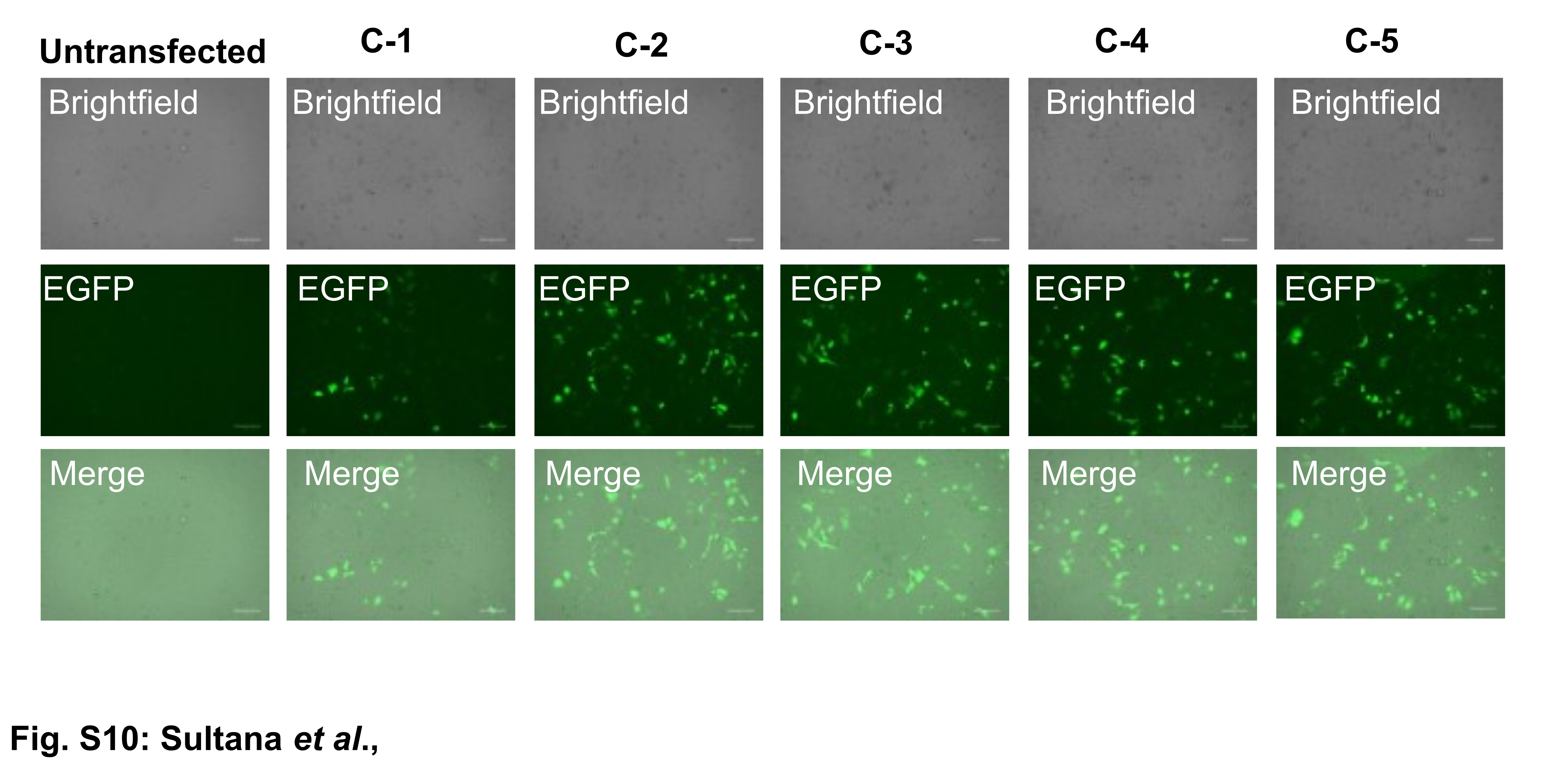

**Supplementary Figure S10:** Transfection of MDA-MB-231 cells with pCMV-EGFP miR-E Notch1 shRNA2 clones (C1-C5)

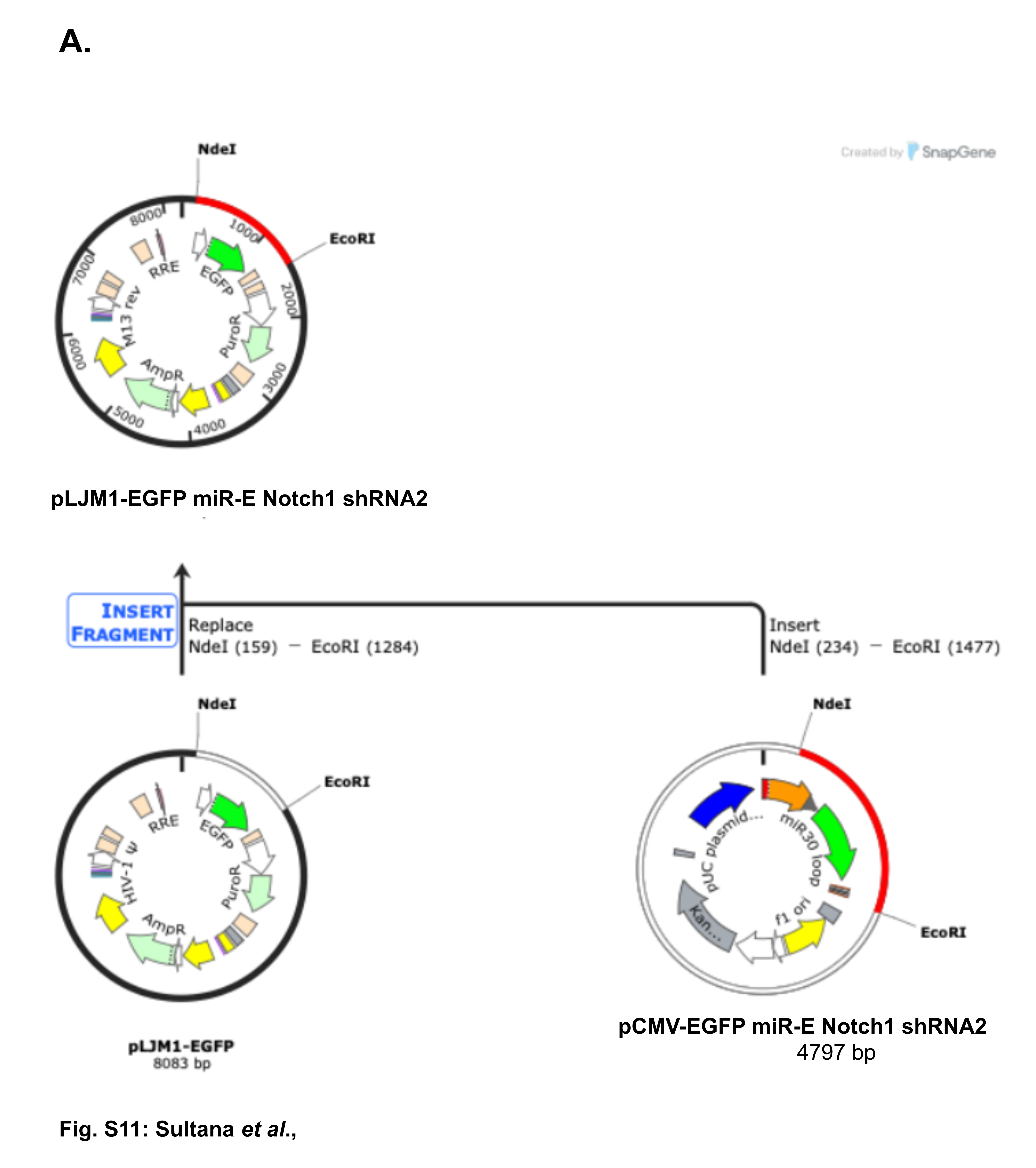

**Supplementary Figure S11:** Schematic representation of the sub-cloning of CMV-EGFP miR-E Notch1 shRNA2 expression cassette in pLJM1-EGFP vector.

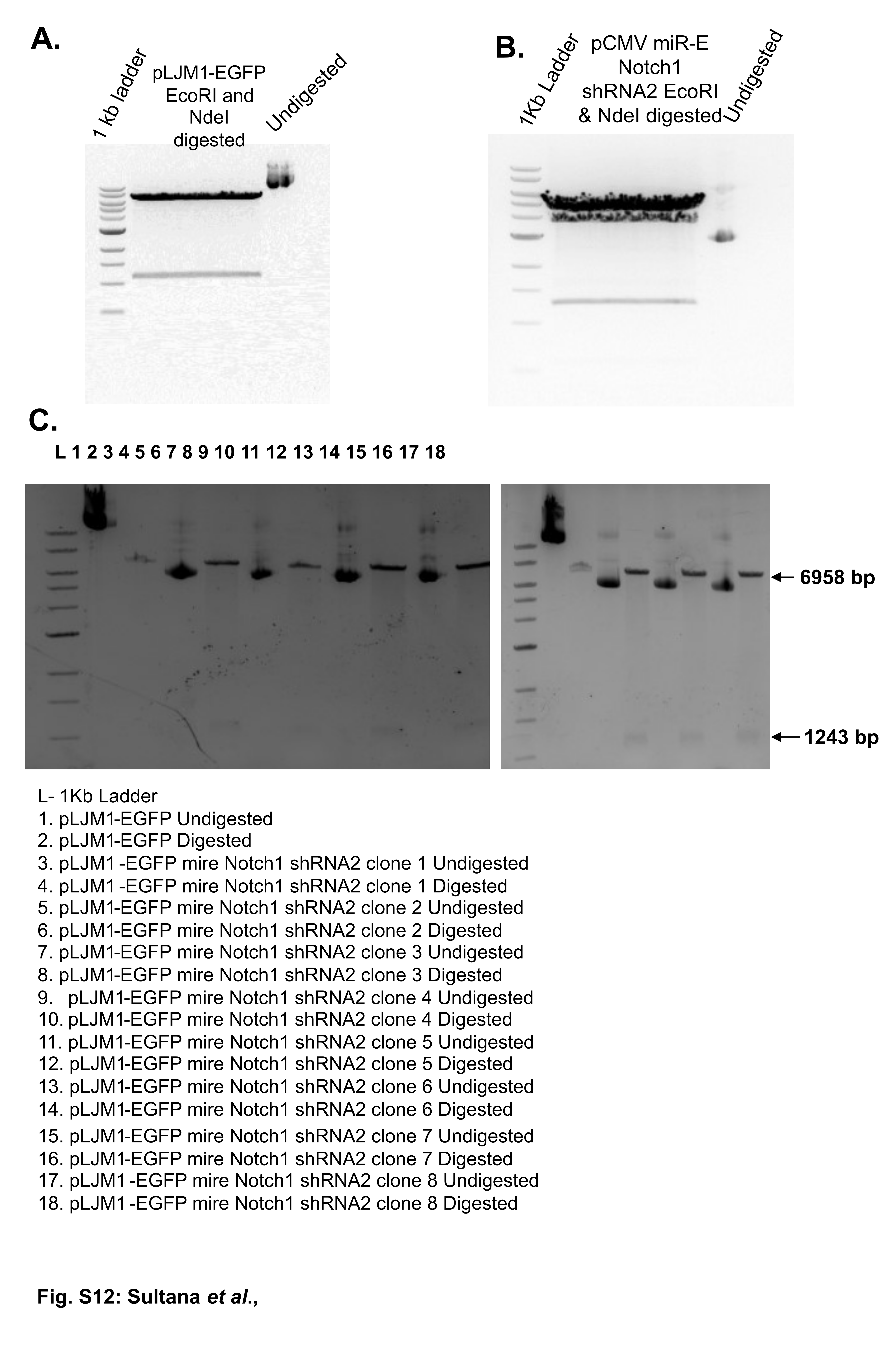

**Supplementary Figure S12:** Sub-cloning of CMV-EGFP miR-E shRNA2 expression cassette in pLJM1-EGFP vector

(A) Restriction digestion of pLJM1-EGFP vector with NdeI and EcoRI enzyme
(B) Restriction digestion of pCMV-EGFP miR-E notch1 shRNA2 with NdeI and EcoRI enzyme.
(C) Restriction mapping of recombinant clones with NheI and BamHI enzyme to confirm successful insertion.

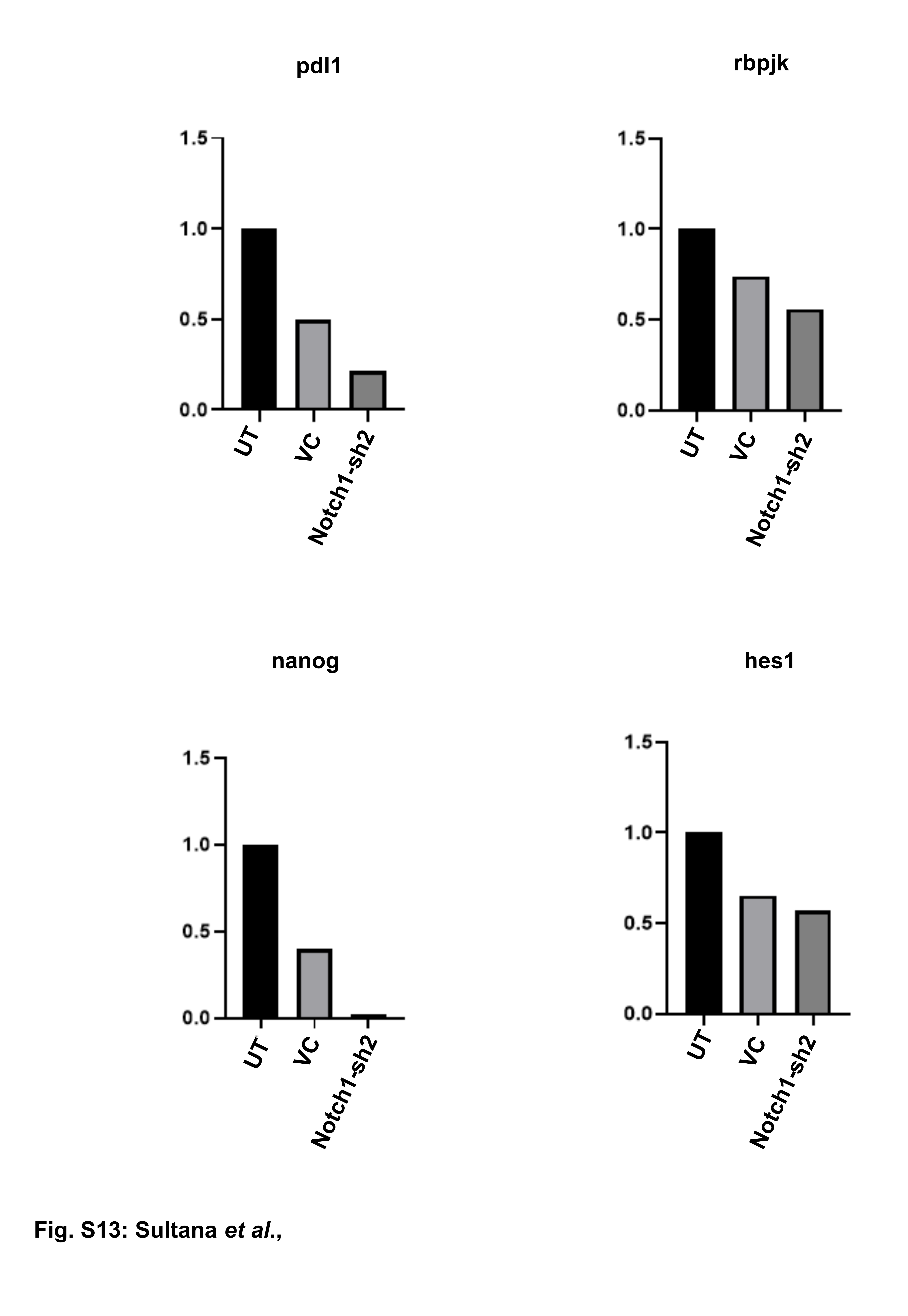

**Supplementary Figure S13:** Quantitative Real time PCR to analyze the level of expression of the genes downstream of Notch1 (NANOG, HES1, PD-L1, and RBPJΚ) in the MDA-MB-231 cells with NOTCH1 knockdown samples as compared vector control (VC) and Untransfected (UT). The level of expression of all 4 downstream targets was found to be downregulated.

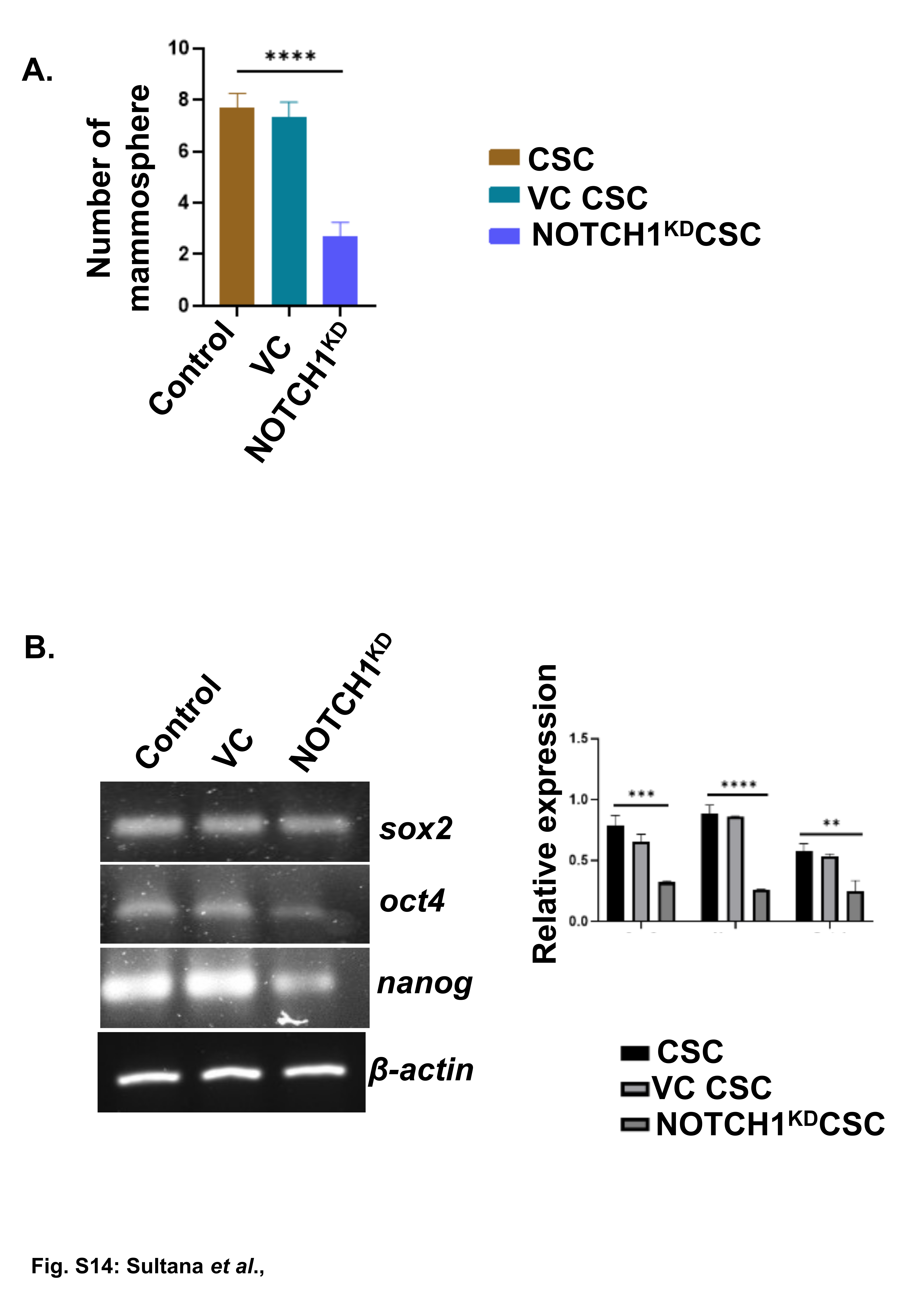

**Supplementary Figure S14:** (A) In bar-graph, mean±SD for mammosphere count is indicated in respective groups of MDAMB-231 cells (Control, Vehicle Control and NOTCH1^KD^). Statistical significance is inferred from two-way ANOVA followed by Tukey’s multiple comparison test (n=3). (B) RT-PCR to analyze the level of expression CSC-fate-regulating genes within CSC-population (*sox2, oct4, nanog*) in the respective groups of MDAMB-231 cells (Control, Vehicle Control and NOTCH1^KD^). Relative fold changes in gene expression across these groups are displayed via bar-graphs (mean± SD); one-way ANOVA analysis followed by Tukey’s multiple-comparison-test (n=5). *p< 0.01, ***p < 0.001, ****p < 0.0001, are indicated.
