## Supplementary Tables T1-T5 for "Breast cancer stem cells mediated CD8^+^ T cell exhaustion among different molecular subtypes of breast cancer regulated via NOTCH1/RBPJ/PD-L1 axis"

**Dr. Saptak Banerjee**

Senior Scientific Officer

Department of Immunoregulation and Immunodiagnostics

Chittaranjan National Cancer Institute

37, S.P. Mukherjee Road

Kolkata – 700026, India

Contact: +91-9717615367

**Supplementary Table T1-**

**Table 1:** Reagent Antibody Details.

| Sl.No | Antibody/Recombinant/Antibody Cocktail kit | Make | Catalogue |
| --- | --- | --- | --- |
| 1. | Lineage Cell Depletion Kit, human | MiltenyiBiotec | 130-092-211 |
| 2. | CD44 MicroBeads, human | MiltenyiBiotec | 130-095-194 |
| 3. | CD24 MicroBead Kit, human | MiltenyiBiotec | 130-095-951 |
| 4. | APC anti-mouse/human CD44 Antibody | Biolegend | 103011  RRID:AB_312962 |
| 5. | FITC anti-mouse/human CD24 Antibody | abcam | AB30350  RRID:AB_726282 |
| 6. | Purified Nanog Antibody | Santa Cruz | sc-374103  RRID:AB_10918255 |
| 7. | Purified Nanog Antibody | GeneTex | GT3312  RRID:AB_11169335 |
| 8. | Purified Nanog Antibody | Cell Signalling Technology | 14955  RRID:AB_2798659 |
| 9. | Purified OCT4 Antibody | R & D Systems | MAB1759  RRID:AB_2167713 |
| 10. | Purified Sox2 Antibody | GeneTex | GTX101507 RRID:AB_2038021 |
| 11. | Human IL-2 Recombinant Protein | PeproTech | AF-200-02  10UG |
| 12. | Human GM-CSF Recombinant Protein | PeproTech | 17831823 |
| 13. | Human IL-4 Recombinant Protein | PeproTech | 200-04-5UG |
| 14. | Biotin anti-human CD8 | Biolegend | 344720  RRID:AB_2075392 |
| 15. | Biotin anti-human CD14 Antibody | Biolegend | 325623  RRID:AB_2074053 |
| 16. | Purified anti-human CD3 | Biolegend | 317301  RRID:AB_571926 |
| 17. | Cy-chrome anti-human CD8 | BD Biosciences | 555368  RRID:AB_395771 |
| 18. | PE- mouse anti-human CD8 | BD Biosciences | 555365  RRID:AB_395768 |
| 19. | Purified anti-human PD-1 (CD279) | Biolegend | 367402  RRID:AB_2565782 |
| 20. | Purified PD-1 antibody | Santa Cruz | sc-10295  RRID:AB_2299183 |
| 21. | PE anti-human CD366 (Tim-3) Antibody | Biolegend | 345005  RRID:AB_1877236 |
| 22. | Brilliant Violet 421™ anti-mouse/human KLRG1 (MAFA) Antibody | Biolegend | 138413  RRID:AB_10918627 |
| 23. | KLRG1 Monoclonal Antibody (13F12F2), Brilliant Violet™ 786. | invitrogen | 417-9488-41  RRID:AB_2937227 |
| 24. | Zombie NIR™ Fixable Viability | Biolegend | 423105 |
| 25. | FITC anti-human CD3 Antibody | Biolegend | 300405  RRID:AB_314059 |
| 26. | PE Mouse Anti-Human Granzyme B | BD Pharmingen | 561142  RRID:AB_10561690 |
| 27. | BV510 Mouse Anti-Human LAG-3 (CD223) | BD Biosciences | 569616  RRID:AB_3685182 |
| 28. | Purified anti-human IFN-γ | ebiosciences | 14-7317-85  RRID:AB_468474 |
| 29. | Purified antibody Pdcd-1L1 | Santa Cruz | sc-518027 |
| 30. | Purified β-actin | Santa Cruz | sc-47778  RRID:AB_626632 |
| 31. | Purified Notch1 antibody | Santa Cruz | sc-376403  RRID:AB_11149738 |
| 32. | Purified Notch1 antibody | Cell signalling Technology | 4380S  RRID:AB_10691684 |
| 33. | Purified Histone1 antibody | Santa Cruz | sc-393358  RRID:AB_3665333 |
| 34. | Delta-like protein 1 goat polycolnal (C-20) | Santa Cruz | sc-8155  RRID:AB_668771 |
| 35. | RBPSUH | Cell signalling Technology | 5313T |
| 36. | Anti-Mouse IgG (Fc specific)–FITC antibody produced in goat | Sigma | F5897  RRID:AB_259677 |
| 37. | Anti-Mouse IgG (whole molecule)–R-Phycoerythrin antibody produced in goat | Sigma | P9287  RRID:AB_261243 |
| 38. | Anti-Rabbit IgG (whole molecule)–FITC antibody produced in goat | Sigma | F9887  RRID:AB_259816 |
| 39. | PerCP-Cy™5.5 Rat Anti-Mouse Ig, κ Light Chain | BD Biosciences | 560668  RRID:AB_1727536 |
| 40. | mouse anti-rabbit IgG PE-Cy7 | Santa Cruz | sc-516721 |
| 41. | goat anti-rat IgG-PE | Santa Cruz | sc-3740  RRID:AB_649013 |
| 42. | bovine anti-goat IgG-PerCP-Cy5.5 | Santa Cruz | sc-45107  RRID:AB_1564055 |
| 43. | PE Goat Anti-Mouse Ig (Multiple Adsorption) | BD Biosciences | 550589  RRID:AB_393768 |
| 44. | goat anti-rabbit IgG-PerCP-Cy5.5 | Santa Cruz | sc-45101  RRID:AB_1564001 |
| 45. | APC-Cy7 Rat Anti-Mouse Ig, κ Light Chain | BD Biosciences | 561353  RRID:AB_10646040 |
| 46. | PE Goat Anti-Mouse Ig (Multiple Adsorption) | BD Biosciences | 550589  RRID:AB_393768 |
| 47. | mouse anti-goat IgG-FITC | Santa Cruz | sc-2356  RRID:AB_628489 |

### Supplementary Table T2-

**Table 2:** List of human patient sample details.

| Sl.No | AGE | Hormone Receptor Status | TNM Stage | FNAC |
| --- | --- | --- | --- | --- |
| 1. | 41 | Luminal A | T4N2M0 | Ductal Carcinoma |
| 2. | 56 | Luminal A | T3N2M1 | Infiltrating Ductal Carcinoma |
| 3. | 45 | Luminal A | T2N1MX | Infiltrating Ductal Carcinoma |
| 4. | 40 | Luminal A | T2N1MX | Infiltrating Ductal Carcinoma |
| 5. | 45 | Luminal A | T4N1M0 | Infiltrating Ductal Carcinoma |
| 6. | 67 | Luminal A | T3N1MX | Invasive Ductal Carcinoma |
| 7. | 60 | Luminal A | T4N1MX | Infiltrating Ductal Carcinoma |
| 8. | 62 | Luminal A | T2N1Mx | Ductal Carcinoma |
| 9. | 23 | Luminal A | T3N1Mx | Ductal Carcinoma |
| 10. | 45 | Luminal A | T2N1Mx | Infiltrating Ductal Carcinoma |
| 11. | 54 | Luminal A | T2N0M0 | Infiltrating Ductal Carcinoma |
| 12. | 52 | Luminal A | T2N0M0 | Infiltrating Ductal Carcinoma |
| 13. | 55 | Luminal A | T4N1Mx | Ductal Carcinoma |
| 14. | 41 | Luminal A | T4N2M0 | Infiltrating Ductal Carcinoma |
| 15. | 58 | Luminal A | T3N2M1 | Infiltrating Ductal Carcinoma |
| 16. | 61 | Luminal A | T2N0M0 | Ductal Carcinoma |
| 17. | 40 | Luminal A | T3N1MX | Ductal Carcinoma |
| 18. | 66 | Luminal A | TCbN0MX | Ductal Carcinoma |
| 19. | 59 | Luminal A | CT4bN1MX | Invasive Ductal Carcinoma |
| 20. | 41 | Luminal A | T2N0MX | Ductal Carcinoma |
| 21. | 47 | Luminal A | T2N1MX | Ductal Carcinoma |
| 22. | 32 | Luminal A | T3N1MX | Ductal Carcinoma |
| 23. | 30 | Luminal B | T2N1MX | Ductal Carcinoma |
| 24. | 40 | Luminal B | T2N1MX | Infiltrating Ductal Carcinoma |
| 25. | 50 | Luminal B | T4N1MX | Ductal Carcinoma |
| 26. | 32 | Luminal B | T2N1Mx | Ductal Carcinoma |
| 27. | 50 | Luminal B | T3N2M1 | Ductal Carcinoma |
| 28. | 65 | Luminal B | T4bN1MX | Infiltrating Ductal Carcinoma |
| 29. | 51 | Luminal B | T2N1MX | Invasive Ductal Carcinoma |
| 30. | 48 | Luminal B | T4N1MX | Infiltrating Ductal Carcinoma |
| 31. | 45 | Luminal B | T2N1MX | Invasive Ductal Carcinoma |
| 32. | 46 | Luminal B | T2N2M0 | Ductal Carcinoma |
| 33. | 51 | Luminal B | T2N1M0 | Ductal Carcinoma |
| 34. | 57 | Luminal B | T2N1MX | High grade Ductal Carcinoma |
| 35. | 53 | Luminal B | T2N0Mx | Infiltrating Ductal Carcinoma |
| 36. | 26 | Luminal B | T2N1MX | Ductal Carcinoma |
| 37. | 60 | Luminal B | T2N1M0 | Ductal Carcinoma |
| 38. | 54 | Luminal B | T2N1MX | Ductal Carcinoma |
| 39. | 35 | Luminal B | T4N1MX | Infiltrating Ductal Carcinoma |
| 40. | 37 | Luminal B | T2N1MX | Ductal Carcinoma |
| 41. | 43 | Her2^+^ | T2N2M0 | Invasive Breast Cancer |
| 42. | 36 | Her2^+^ | T3N2M0 | Invasive Ductal Carcinoma |
| 43. | 40 | Her2^+^ | T4bN1Mx | Invasive Ductal Carcinoma |
| 44. | 58 | Her2^+^ | T2N2M0 | Invasive Breast Cancer |
| 45 | 59 | Her2^+^ | T4bN1Mx | Invasive Breast Cancer |
| 46 | 45 | Her2^+^ | T2N1Mx | Invasive Breast Carcinoma |
| 47 | 47 | Her2^+^ | T2N1Mx | Invasive Breast Carcinoma |
| 48 | 25 | Her2^+^ | T3N1Mx | Infiltrating Ductal Carcinoma |
| 49 | 51 | Her2^+^ | T3N1Mx | Ductal Carcinoma |
| 50 | 57 | Her2^+^ | T2N1Mx | Ductal Carcinoma |
| 51 | 52 | Her2^+^ | T2N1Mx | Infiltrating Ductal Carcinoma |
| 52 | 62 | Her2^+^ | T4bN1Mx | Invasive Breast Carcinoma |
| 53 | 44 | Her2^+^ | T3N1Mx | Invasive Ductal Carcinoma |
| 54 | 50 | Her2^+^ | T2N1Mx | Invasive Breast Carcinoma |
| 55 | 47 | Her2^+^ | T2N0M0 | Ductal Carcinoma |
| 56 | 56 | Her2^+^ | T2N1M0 | Invasive Ductal Carcinoma |
| 57 | 56 | Her2^+^ | T2N1Mx | Invasive Breast Carcinoma |
| 58 | 79 | Her2^+^ | T4bN1M0 | Ductal Carcinoma |
| 59 | 61 | Her2^+^ | T2N1M0 | Ductal Carcinoma |
| 60 | 52 | Her2^+^ | T4bN1M0 | Invasive Ductal Carcinoma |
| 61 | 55 | Her2^+^ | T4bN1M0 | Invasive Ductal Carcinoma |
| 62 | 53 | Her2^+^ | T3N1M0 | Invasive Ductal Carcinoma |
| 63 | 55 | Her2^+^ | T2N1M0 | Invasive Ductal Carcinoma |
| 64 | 63 | TNBC | T3N1Mx | Invasive Ductal Carcinoma |
| 65 | 55 | TNBC | T3N1Mx | Infiltrating Ductal Carcinoma |
| 66 | 30 | TNBC | T2N1M0 | Invasive Ductal Carcinoma |
| 67 | 56 | TNBC | T4N1Mx | Invasive Ductal Carcinoma |
| 68 | 50 | TNBC | T3N1Mx | Invasive Mammary Carcinoma |
| 69 | 52 | TNBC | T2N2Mx | Infiltrating Ductal Carcinoma |
| 70 | 58 | TNBC | T2N0M0 | Invasive Mammary Carcinoma |
| 71 | 36 | TNBC | T4N2M1 | Invasive Ductal Carcinoma |
| 72 | 51 | TNBC | T3N2M1 | Invasive Ductal Carcinoma |
| 73 | 50 | TNBC | T3N2M1 | Invasive Ductal Carcinoma |
| 74 | 58 | TNBC | T4N2M1 | Invasive Ductal Carcinoma |
| 75 | 63 | TNBC | T2N2M1 | Invasive Ductal Carcinoma |
| 76 | 51 | TNBC | T3N1M0 | Not given |
| 77 | 52 | TNBC | T3N1M0 | Not given |
| 78 | 52 | TNBC | T3N1M0 | Infiltrating Ductal Carcinoma |
| 79 | 34 | TNBC | T4bN3Mx | Invasive Ductal Carcinoma |
| 80 | 65 | TNBC | T4bN1M0 | Invasive Breast Carcinoma |
| 81 | 63 | TNBC | T4N3M1 | Invasive Ductal Carcinoma |
| 82 | 41 | TNBC | T4bN3Mx | Invasive Breast Carcinoma |
| 83 | 58 | TNBC | T3N2M0 | Invasive Breast Carcinoma |
| 84 | 52 | TNBC | T4N3Mx | Invasive Ductal Carcinoma |
| 85 | 56 | TNBC | T4N1Mx | Invasive Ductal Carcinoma |
| 86 | 55 | TNBC | T3N1Mx | Invasive Ductal Carcinoma |
| 87 | 52 | TNBC | T2N2Mx | Ductal Carcinoma |
| 88 | 47 | TNBC | T3N2M0 | Infiltrating Ductal Carcinoma |
| 89 | 64 | TNBC | T4N3M1 | Invasive Ductal Carcinoma |
| 90 | 58 | TNBC | T4N2M0 | Infiltrating Ductal Carcinoma |
| 91 | 54 | TNBC | T4N3M1 | Infiltrating Ductal Carcinoma |
| 92 | 52 | TNBC | T4N3M0 | Infiltrating Ductal Carcinoma |
| 93 | 61 | TNBC | T3N2M1 | Invasive Ductal Carcinoma |
| 94 | 63 | TNBC | T4N3M1 | Invasive Breast Carcinoma |
| 95 | 52 | TNBC | T3N1AMx | Invasive Ductal Carcinoma |

**Supplementary Table T3-**

**Table 3: Primer Details used in the study.**

| SL.No | Primer Name | Forward primer sequence 5’- 3’ | Reverse primer sequence |
| --- | --- | --- | --- |
| 1. | Human *β-actin* | 5’-GACATTAAGGAGAAGCTGTG-3’ | 5’-GAGTTGAAGGTAGTTTCGTG-3’ |
| 2. | Human *cd44* | 5’-AATACAGAACGAATCCTGAA -3’ | 5’-GATGGGGTGTACAGTAGAAA -3’ |
| 3. | Human *sox2* | 5’-CAGTACAACTCCATGACCAG -3’ | 5’-AGTGGGAGGAAGAGGTAAC -3’ |
| 4. | Human *oct4(2)* | 5’-GGGTTCTATTTGGGAAGG -3’ | 5’-GATACTGGTTCGCTTTCT -3’ |
| 5. | Human  *nanog* | 5’-AAGAATAGCAATGGTGTGAC-3’ | 5’-TTATAGAAGGGACTGTTCCA -3’ |
| 6. | Human  *notch3* | 5’-GGCGCTTCCTGGACAATCAT -3’ | 5’-CATCCCCAAACCACACTCGT -3’ |
| 7. | Human *notch2* | 5’-CTGTGTGTTTGCCGTAGTGC-3’ | 5’-CACTTGTCCACAGCTGCTCT-3’ |
| 8. | Human *notch1* | 5’-GAACTGTGAGGAAAATATCG-3’ | 5’-GACACACACGCAGTTGTAG-3’ |
| 10. | Human *notch4* | 5’-AGAAAGACTCCACCTTTCAC-3’ | 5’-GTCTCACACTCATCCACATC-3’ |
| 11. | Human *hes1* | 5’-AGCACAGAAAGTCATCAAAG-3’ | 5’-TTCACTGTCATTTCCAGAAT -3’ |
| 12. | Human *hey1* | 5’-AAAAGACGGAGAGGAATAAT-3’ | 5’- CAAACTCCGATAGTCCATAG-3’ |
| 13. | Human *eomes* | 5’-TGTTCTCTGAAGATCAGCTC -3’ | 5’-AATGTCCTCACACTTTATGG -3’ |
| 14. | Human *ifnγ* | 5’GGACCCATATGTAAAAGAAGCAGA -3’ | 5’-TGTCACTCTCCTCTTTCCAATTCT -3’ |
| 15. | Human *tox* | 5’-CTGACTACCATTAACCAGTC -3’ | 5’-CTCTCCACCATTGATCTTAG -3’ |
| 16. | Human *pdcd1* | 5’-CTCAGGGTGACAGAGAGAA -3’ | 5’-CCATAGTCCACAGAGAACAC-3’ |
| 17. | Human *klrg1* | 5’- TGCCTAGCCAGAGACTCACA-3’ | 5’-CACCGCATGTCTGCACAAAG -3’ |
| 18. | Human *tim3* | 5’-ACATCCAGATACTGGCTAAA -3’ | 5’-TGACCAACTTCAGGTTAAAT -3’ |
| 19. | Human *batf* | 5’-CTGGGCAGCAGACAAATCCT-3’ | 5’-CACAACCAAGGTGCTAGCCA-3’ |
| 20. | Human  *rbpj-nanog* | 5’-CAGACCTGGGAAGAAGCTAAAGA -3’ | 5’-GGTACTTGTTTTCAGCACCTACC -3’ |
| 21. | Human  *rbpj-pdl1* | 5’-TGGCAGAATATCAGGGACCCT -3’ | 5’-CGTGGATTCTGTGACTTCCTCA -3’ |

**Supplementary Table T4-**

**Table. 4:** List of shRNA sequences used in the study.

| **Sl.No.** | **shRNA** | **Sequence** |
| --- | --- | --- |
| 1 | Notch1_shRNA1 | 5’ CCGGGACATCACGGATCATAT 3’ |
| 2 | Notch1_shRNA2 | 5’ CTGAACCAGGGCACGTGTATT 3’ |
| 3 | Notch1_shRNA3 | 5’ GGACAAGATCGATGGCTACGA 3’ |
| 4 | Notch1_shRNA4 | 5’ GCTCACGCTGACGGAGTACAA 3’ |

**Supplementary Table T5-**

**Table. 5: List of primers used in the study for knockdown experiments.**

| **Target** | **Primer name** | **Sequence** |
| --- | --- | --- |
| Nanog | Nanog RT_FP | 5’ CAATGGTGTGACGCAGAAGG 3’ |
|  | Nanog RT_RP | 5’ TGCTCCAGGACTGGATGTTC 3’ |
| RBPJκ | RBPJk RT_FP | 5’ CTTATTCTCTCGGCACCCC 3’ |
|  | RBPJk RT_RP | 5’ ACCAAATTTCCGTAGAGTCTTG 3’ |
| PD-L1 | PDL1_RT_FP | 5’ ATAGGCCAATGTGGTCTGGG 3’ |
|  | PDL1_RT_RP | 5’ ATGGCAAAGGCAAATCAGGAAT 3’ |
| Hes1 | Hes1_RT_FP | 5’ ACACGACACCGGATAAACCA 3’ |
|  | Hes1_RT_RP | 5’ GGAATGCCGCGAGCTATCTT 3’ |
| Notch1 | Notch1_RT_FP | 5’ AGACTATGCCTGCAGCTGTG 3’ |
|  | Notch1_RT_RP | 5’ GGCACGATTTCCCTGACCA 3’ |
